# PTPRZ1-targeting RNA CAR-T cells exert antigen-specific and bystander antitumor activity in glioblastoma

**DOI:** 10.1101/2023.12.23.573190

**Authors:** Darel Martinez Bedoya, Eliana Marinari, Suzel Davanture, Luis Cantero Castillo, Sarah Erraiss, Millicent Dockerill, Sofia Barluenga Badiola, Nicolas Winssinger, Karl Schaller, Philippe Bijlenga, Shahan Momjian, Philippe Hammel, Pierre Cosson, Paul R. Walker, Valérie Dutoit, Denis Migliorini

## Abstract

The great success of chimeric antigen receptor (CAR)-T cell therapy in B-cell malignancies has prompted its translation to solid tumors. In the case of glioblastoma (GBM), clinical trials have shown modest efficacy, but anti-GBM CAR-T cells are being intensely developed. In this study, we selected PTPRZ1 as an attractive new target for GBM treatment. We isolated six anti-human PTPRZ1 scFv from a human phage display library and produced 2^nd^ generation CAR-T cells in an RNA format. Patient-derived GBM PTPRZ1-knock-in cell lines were used to select the CAR construct (471_28z), which showed high cytotoxicity while consistently displaying high CAR expression. CAR-T cells incorporating 471_28z were able to release IFN-γ, IL-2, TNF-α, Granzyme B, IL-17A, IL-6, and soluble FasL, and displayed low tonic signaling. Additionally, they maintained an effector memory phenotype after *in vitro* killing. Importantly, 471_28z CAR-T cells displayed strong bystander killing against PTPRZ1-negative cell lines after pre-activation by PTPRZ1-positive tumor cells, but did not kill antigen-negative non-tumor cells. In an orthotopic xenograft tumor model using NSG mice, a single dose of anti-PTPRZ1 CAR-T cells significantly delayed tumor growth. Taken together, these results validate the use of PTPRZ1 as a new GBM target and prompt the use of anti-PTPRZ1 CAR-T cells for clinical translation.

## Introduction

Glioblastoma (GBM) is the most frequent primary tumor originating in the brain, and despite available treatments, its recurrence is universal. In the last few decades, researchers have identified GBM antigens that allow for the development of immunotherapies (1,2). Despite some promising results with multi-peptide vaccines (3,4) or neoadjuvant immune checkpoint blockers (5,6), most clinical trials in GBM have been disappointing.

Adoptive cell transfer with chimeric antigen receptor (CAR)-T cells has been a breakthrough therapy against cancer in recent years, with unprecedented success in hematological malignancies (7,8). Attempts have been made to translate the success of CAR-T cells to GBM, and clinical trials have shown an encouraging safety profile and occasional clinical benefit but still have an insufficient impact on overall survival (9,10). CAR-T cell therapy remains one of the most promising alternatives for GBM, but it is necessary to increase its efficacy, which is currently limited by tumor heterogeneity and an immunosuppressive tumor microenvironment, by targeting more relevant antigens and improving CAR-T cell functionality, among others (11).

Through peptidomics analysis, our group identified PTPRZ1 as an immunogenic and tumor-relevant GBM-associated antigen and used it as part of the IMA950 multi-peptide vaccine in phase I/II clinical trials (4,12,13). PTPRZ1 (Protein tyrosine phosphatase receptor type Z, also named RPTPβ) is a membrane-linked protein with tyrosine phosphatase activity in its cytoplasmic domain and is a key element in regulating the differentiation of oligodendrocyte precursors (14). PTPRZ1 is highly expressed in gliomas, especially GBM, and plays a role in tumor growth, invasion, radioresistance, and generation and maintenance of the immunosuppressive tumor microenvironment (15,16). In the absence of its ligands, monomeric PTPRZ1 is constitutively active and dephosphorylates its targets (such as ALK, c-Src, β-catenin, or MAGI proteins), leading to the inhibition of their respective pathways. Binding of PTPRZ1 to its ligands, such as pleiotropin, midkine, or IL-34, causes receptor homodimerization and inhibition of phosphatase activity, allowing increased phosphorylation of its targets and activation of tumorigenic pathways (17). Interestingly, PTPRZ1 has been described as a relevant marker of glioma stem cells, a population considered responsible for chemo- and radio-resistance, and therefore for tumor recurrence (18,19). In recent years, different means of inhibiting its activity have been described using RNA interference, small molecules, or antibody-directed nanoparticles (16,20), but although some promising preclinical results have been obtained, no therapy targeting PTPRZ1 has reached the clinical stage.

We previously detected PTPRZ1 protein expression in the majority of patients with newly diagnosed and recurrent GBM (4,12), and we used this antigen to develop GBM-targeting CAR-T cells. We chose to design CAR-T cells in mRNA format, considering the advantages of reduced manufacturing time and costs, improved safety, and flexibility (21,22). Transient CAR expression of mRNA-based CAR-T cells could mitigate potential toxicity once infused into the patient, allowing repetitive infusions. PTPRZ1-specific single-chain variable fragments (scFv) were isolated and tested *in vitro* using PTPRZ1-expressing patient-derived GBM lines and *in vivo* in NSG mice, resulting in the selection of one scFv with consistently high CAR expression and high cytotoxic activity, as well as high cytokine and effector protein secretion. Our results validated the use of PTPRZ1-trageting CAR-T cells as an attractive new option for treating GBM.

## Materials and Methods

### Gene expression data analysis

Bulk RNA-seq data from TCGA and Rembrandt were analyzed and plotted using GlioVis (23). For healthy tissues, GTEx RNA-seq data were used (GTEx Analysis Release V8, dbGaP Accession phs000424.v8.p2). Single-cell data from Neftel et al. (24) were analyzed, and figures were generated using the single-cell portal (https://singlecell.broadinstitute.org). Single-cell data from Yuan et al. (2018) (geo_accession"GSE103224")(25) were downloaded and analyzed. Seurat v3 (26,27) was used for all subsequent analyses. We constructed a Seurat object using a feature barcode matrix for each sample. A series of quality filters were applied to the data to remove low-quality cell barcodes: possible debris with too few genes expressed (< 300), possibly more than one cell with too many genes expressed (> 6,000-10,000 according to the sample), possible dead cells, or a sign of cellular stress and apoptosis with an extremely high proportion of mitochondrial gene expression over the total transcript counts (> 20%). Each sample was scaled and normalized using Seurat’s NormalizeData’ and ‘ScaleData’ functions, respectively. We then merged all the samples and repeated the same scaling and normalization method. All cells in the merged Seurat object were integrated using ‘IntegrateData’ and the top 30 PCA dimensions were then clustered via Seurat’s FindNeighbors’ and ‘FindClusters’ (with parameters: resolution = 0.5) functions. The resulting merged and normalized matrices were used for subsequent analyses. Differentially expressed genes within immune and tumor cells were identified using FindMarkers function by comparing cells belonging to one subtype to the rest. Wilcoxon signed-rank test was used. log2FC > 0.25 and FDR < 0.05, were used to filter DEGs. Clusters of tumor cells were annotated as described in this study.

### Cell lines and reagents

The human GBM cell lines Ge518 and Ge1302 were generated in our laboratory from brain tumor resections. They were cultured as adherent lines in Dulbecco’s modified Eagle’s medium supplemented with high glucose (4.5 g/L) and sodium pyruvate (DMEM, Gibco, 31966-021), and enriched with 8% human serum (Sigma, H4522), 10 mM HEPES (Gibco, 15630-056), MEM Amino Acids Solution (Gibco, 11140-050) and 1% Penicillin-Streptomycin (Gibco, 15140-122). Cells were cultured at 37°C with 5% CO_2_ and were regularly tested for mycoplasma contamination using the MycoAlert® PLUS Mycoplasma Detection Kit (Lonza, LT07-703), and authenticated by Microsynth. Jurkat triple reporter cells, containing NF-κB, NFAT, and AP-1 T cell response elements linked to CFP, eGFP, and mCherry, respectively, were kindly donated by Peter Steinberger’s lab (Medical University of Vienna, Vienna, Austria) to evaluate CAR tonic signaling (28).

### ScFv isolation

Human PTPRZ1 (Unitprot P23471)-carbonic anhydrase and fibronectin type III extracellular domains were cloned into the Peak8del vector (29) as a biotinylated chimeric protein comprising a human IgG1 Fc portion. The chimeric hIgG1 Fc-PTPRZ1 protein was produced by episomal vector expression in HEK-derived PeakRapid cells (ATCC, CRL-2828, RRID:CVCL_3772). To confirm the domain sequences, samples were digested, and peptides were analyzed by nanoLC-MSMS using an easy nLC1200 (Thermo Fisher Scientific) coupled with an Orbitrap Fusion Lumos mass spectrometer (Thermo Fisher Scientific). Database searches were performed with Mascot (Matrix Science, RRID:SCR_014322), and data were analyzed and validated with Scaffold (Proteome Software, RRID:SCR_014321) with 95% probability for both peptides and proteins, and at least two unique peptides per protein. A phage display library of human single-chain variable fragments (scFv) (30) was screened with the proteins produced to isolate scFvs capable of recognizing PTPRZ1 extracellular domains. Binding of the isolated scFv-Rabbit Fc (ABCD_RB469, ABCD_RB470, ABCD_RB471, ABCD_RB473, ABCD_RB474 and ABCD_RB476 as referenced in https://web.expasy.org/abcd/) to PRPRZ1 extracellular domains was measured by ELISA.

### Surface plasmon resonance (SPR) measurements

SPR experiments were performed using a Biacore T200 instrument (GE Healthcare) at 25°C in PBS-P+ buffer (10X stock, Cytiva Life Sciences, 28995084). The scFvs were captured on a Protein G Sensor Chip (Cytiva Life Sciences, 29179316) by flowing a solution of 0.5 μg/mL of the compound in PBS-P+ for 120 seconds at a flow rate of 10 μL/min. The chip was allowed to stabilize for 80 s prior to kinetic measurements. The latter consisted of injections (association 250 s, dissociation 500 s, flow rate: 30 μL/min) of decreasing concentrations of the PTPRZ1 CA (Cusabio Technology LLC, CSB-YP019068HU) or CAII (Sigma Aldrich, C2522) proteins (2-fold cascade dilutions from the starting concentration of 800 nM). The chip was regenerated between cycles by one injection of the regeneration solution (10 mM glycine-HCl at pH 1.5, Cytiva Life Sciences, BR100354) for 30 s at a flow rate of 20 μL/min. Binding was measured as resonance units over time after blank subtraction, and the data were interpreted using Biacore T200 software, version 3.2 (Cytiva Life Sciences, RRID:SCR_019718). *K*_D_ values were calculated based on steady-state affinity analysis. This model (Equation 1) calculates the equilibrium dissociation constant KD for a 1:1 interaction from a plot of steady-state binding levels (R_eq_) against analyte concentration (C). The equation includes an offset term that represents the response at zero analyte concentration. The response at steady state was calculated 4 s prior to the injection stop, with a window of 5 s.

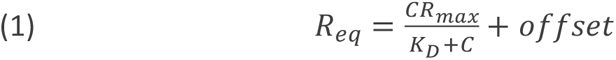

### CAR design and cloning

The gene coding for each of the six anti-PTPRZ1 scFv genes was synthesized as a gene-encoding plasmid using GeneArt (Thermo Fisher Scientific). In all cases, the restriction enzymes BamHI-HF (New England Biolabs, R3136) and BspEI (New England Biolabs, R0540) were used to clone the scFv fragment in a vector optimized for mRNA expression (pDA) (31). To complete the CAR construct, the pDA vector contained (1) human CD8 hinge and transmembrane segments followed by a human 4-1BB intracellular domain, (2) human IgG4 hinge followed by a human CD28 transmembrane domain and intracellular domain, or (3) human IgG1 hinge followed by human CD28 transmembrane and intracellular domains. All constructs started with a human CD8a leader sequence, and a human CD3ζ intracellular domain was present at the end of the CAR structure. Plasmids containing the full CAR constructs were used to transform One Shot Top10 chemically competent cells (Invitrogen, C404003), and bacteria were grown overnight (ON) at 37 °C with shaking. DNA was extracted using the GeneJET Plasmid Maxiprep Kit (Thermo Fisher Scientific, K0492).

### mRNA production

Plasmids with CAR constructs were linearized by digestion with the restriction enzyme SalI (New England Biolabs). mRNA was produced using the mMESSAGE mMACHINE™ T7 ULTRA Transcription Kit ( AMB13455; Invitrogen) and later cleaned with the MEGAclear Transcription Clean-Up Kit ( AM1908; Invitrogen). mRNA quality was checked by 1% agarose electrophoresis, and then mRNA was resuspended at 3 μg/μL and frozen at −80°C.

### RNA CAR-T cell generation

Human blood or buffy coats (Interregionale Blutspende SRK AG, Bern, Switzerland) were used as the starting materials for T cell isolation. T cells were obtained from whole blood using the RosetteSep™ Human T Cell Enrichment Cocktail (StemCell Technologies, 15061), followed by density gradient separation using Ficoll® Paque Plus (Cytiva Life Sciences, 17144003). From buffy coats, human peripheral blood mononuclear cells (PBMC) were obtained through density gradient separation using Ficoll® Paque Plus, and T cells were purified using the EasySep™ Human T Cell Isolation Kit (StemCell Technologies, 17951). Once purified, T cells were diluted to 1×10^6^ cells/mL and activated with a 1:1 ratio of anti-CD3 and anti-CD28 Dynabeads (Gibco, 11141D). After 48h of incubation, beads were magnetically removed and activated T cells were washed with MaxCyte Electroporation buffer (Cytiva Life Sciences, EPB1). T cells were electroporated with 10 μg of mRNA for 3×10^6^ cells using ExPERT GTx™ (MaxCyte). Following electroporation, T cells were incubated at 37°C with 5% CO_2_ in RPMI1640 medium supplemented with 10% fetal bovine serum (FBS, Sigma, F7524-500mL), 1 mM sodium pyruvate (Gibco, 11360-070), 10 mM HEPES (Gibco, 15630-056) and 1% penicillin-streptomycin (Gibco, 15140-122). After the first 2h of incubation, the medium was supplemented with 30 IU/mL recombinant human interleukin 2 (IL-2) (Proleukin, Roche). Electroporated T cells were kept in culture ON and then used directly or frozen.

### PTPRZ1-KI (knock-in) and KO (knock-out) cell generation

PTPRZ1-KI lines were generated from Ge518 and Ge1302 cell lines by transduction of wild-type lines with a lentivirus containing the peptide leader, α-carbonic anhydrase, and fibronectin type III extracellular domains (aa 1–419), followed by the transmembrane domain (aa 1636– 1671) of human PTPRZ1 (gene synthesized by GeneArt, Thermo Fisher Scientific). Resistance to puromycin (InvivoGen, ant-pr-1) was used as a selection marker. Lentivirus was produced using a 3^rd^ generation vector, pCDHv6_PTPRZ1 (derived from pCDH-MSCV-MCS-EF1-GFP-T2A-Puro, System Biosciences, CD713B-1), with packaging plasmids pMDLg/pRRE (RRID:Addgene_12251), pRSV-Rev (RRID:Addgene_12253), and pMD2.G (RRID:Addgene_12259) to express the VSV-G envelope. The Ge518_PTPRZ1-KO cell line was generated through CRISPR/Cas9 editing using sgRNA 5’-UUUUUGAUUCAGUGCUCCUA-3’ designed using the CRISPR Design Tool (https://www.synthego.com/products/bioinformatics/crispr-design-tool). Ge518 (wild-type) cells were electroporated with 112 pmol sgRNA and 2 μg TrueCut Cas9 Protein v2 (Thermo Fisher Scientific, A36498) using a 4D-Nucleofector (Lonza). After 72h of culture, the cells were cloned by seeding 1 cells/well in a 96-well plate. Genomic DNA was extracted using the PureLink™ Genomic DNA Mini Kit (Thermo Fisher Scientific, K182000) and the PTPRZ1 sequence was checked by PCR using AccuPrime™ Pfx SuperMix (Thermo Fisher Scientific, 12344040) with the forward 5’-AGTGGCATGGTTGTTCTACAGACC-3’ and reverse 5’-GAAACTAGTCTAAACATACTCCAGGTCTG-3’ primers (Microsynth). The efficiency of PTPRZ1-KO was measured by comparing the sequence of clones with that of wild-type cells using the ICE CRISPR Analysis Tool (https://www.synthego.com/products/bioinformatics/crispr-analysis). Cell lines were transduced to express GFP, mCherry, or luciferase using lentiviruses, and positive cells were selected using 2 μg/mL puromycin (InvivoGen, ant-pr-1) for GFP and mCherry or 1 mg/mL G-418 solution (Roche, 04727878001) for luciferase. Lentiviruses used to produce GFP, mCherry, or luciferase used the same packaging and envelope plasmids as described previously with the respective 3^rd^ generation plasmids: pCDH-MSCV-MCS-EF1-GFP-T2A-Puro (System Biosciences, CD713B-1), pLenti-mCherry (derived from pLenti-III-EF1a, Applied Biological Materials Inc., LV043), and Lenti-luciferase-P2A-Neo (RRID:Addgene_105621).

### Flow cytometry analysis

CAR surface expression was measured using biotinylated goat anti-human IgG F(ab’)₂ (Jackson Immunoresearch, 109-066-097) followed by streptavidin conjugated to PE (BD Biosciences, 554061) or APC (BD Biosciences, 554067). The LIVE/DEAD™ Fixable Violet Dead Cell Stain Kit (Invitrogen, L34964) was used to discriminate between live and dead cells. To detect PTPRZ1 expression in KI tumor cells, isolated anti-PTPRZ1 scFvs conjugated to rabbit Fc were used, followed by goat anti-rabbit IgG^AF488^ (Invitrogen, A11008). For phenotypic characterization, the following antibodies were used: anti-CD3^FITC^ (BioLegend, 317306), anti-CD8^BV421^ (BioLegend, 344748), anti-CD45RO^PE-Cy7^ (Biolegend, 304230), anti-CD62L^PerCP-Cy5.5^ (Biolegend, 304824), anti-PD-1^APC-Cy7^ (BioLegend, 367416), anti-TIM3^PE-CF594^ (BD Biosciences, 565560), anti-CD127^PE^ (BD Biosciences, 557938), and LIVE/DEAD™ Fixable Yellow Dead Cell Stain Kit (Invitrogen, L34967). Isotype controls used were anti-PD-1 (BioLegend, 400128), anti-TIM3 (BD Biosciences, 562292), and anti-CD127 (BD Biosciences, 554680). Cell acquisition was performed using a BD FACSymphony™ (BD Biosciences). All data were analyzed using FlowJo 10 (Tree Star, RRID:SCR_008520). To evaluate cytokine secretion in the supernatants, we used the LEGENDplexTM Human CD8/NK Panel (13-plex) (Biolegend) following the manufacturer’s instructions, and the data were analyzed using the LEGENDplex™ Data Analysis Software Suite (Qognit). The effector/memory T cell subsets were defined as follows: central memory (T_CM_), CD45RO^+^ CD62L^+^; effector memory (T_EM_), CD45RO^+^ CD62L^-^; effector (T_EFF_), CD45RO^-^ CD62L^-^ and naïve (T_N_), CD45RO^-^ CD62L^+^.

### Flow cytometry-based killing assay

Target tumor cells were stained using the CellTrace™ Far Red Cell Proliferation Kit (Invitrogen, C34564) and 25’000 cells per well were seeded in a 96-well flat-bottom plate. The tumor cells were incubated for 2h at 37°C with 5% CO_2_ to allow them to adhere to the plate. Effector anti-PTPRZ1 RNA CAR-T cells or control mock electroporated (Mock EP) T-cells were added to target cells at different effector-to-target (E:T) ratios. After 72h of co-incubation, cells were collected and tumor cell death was measured by flow cytometry (BD FACSymphony™, BD Biosciences) using a LIVE/DEAD™ Fixable Violet Dead Cell Stain Kit (Invitrogen, L34964). Dead tumor cells were identified as LIVE/DEAD^+^ FarRed^+^ cells. The following formula was used to calculate the percentage of specific lysis:

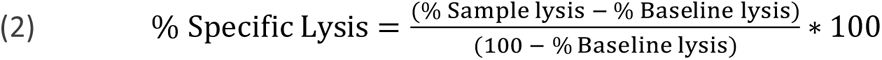

where “% Sample lysis” corresponds to lysis in the presence of Mock EP or CAR-T cells and “% Baseline lysis” corresponds to lysis in the absence of T cells (tumor cells alone).

### Incucyte-based tumor growth inhibition assay

Target tumor cells expressing a fluorophore (GFP or mCherry) were seeded in a 96-well plate at a concentration of 5’000 cells/well for Ge518, for both KI and KO variants, and 15’000 cells/well for Ge1302_PTPRZ1-KI. Tumor cells were incubated for 2h at 37°C with 5% CO_2_ to allow adherence to the plate surface. Effector anti-PTPRZ1 RNA CAR-T cells or control Mock EP T cells were added to target cells at different E:T ratios. Tumor cell growth was monitored for up to 120h by real-time imaging, measuring the total fluorophore area per well every 2h h using an Incucyte® S3 Live-Cell Analysis System (Sartorius). The data from each well were normalized to the first measurement (0h).

### IFN-γ detection by ELISA

IFN-γ secreted in the supernatant after 72h of co-culture of tumor and effector T cells was measured using the Human IFN-gamma DuoSet ELISA kit (R&D Systems, DY285B) following the manufacturer’s instructions. Absorbance of the ELISA plates at 450 and 540 nm was measured using a Spark® Multimode Microplate Reader (Tecan).

### Evaluation of bystander killing through soluble mediators

Ge518_PTPRZ1-KO cells were stained using the CellTrace™ Yellow Cell Proliferation Kit (Invitrogen, C34573), and Ge518_PTPRZ1-KI cells were stained using the CellTrace™ Far Red Cell Proliferation Kit (Invitrogen, C34564). Using Polycarbonate Cell Culture Inserts (0.4 µm pore) in a 6-well plate (Nunc, 140660), 50’000 Ge518_PTPRZ1-KO cells were seeded in the bottom of the well and 50’000 Ge518_PTPRZ1-KI cells were seeded at the top of the insert. At 24h, anti-PTPRZ1 RNA 471_28z CAR-T cells or Mock EP control T cells were added to the top of the transwell at an 5:1 E:T ratio. After 72h, cells on the bottom of the wells were collected, and cell death was evaluated by flow cytometry using a LIVE/DEAD™ Fixable Violet Dead Cell Stain Kit (Invitrogen, L34964).

### Evaluation of bystander killing on non-tumoral human macrophages

Human PBMC were obtained from buffy coats by density gradient separation using Ficoll® Paque Plus, and CD14^+^ cells were purified using the EasySep™ Human CD14 Positive Selection Kit II (StemCell Technologies, 17858). A total of 1.5×10^6^/well CD14^+^ monocytes were seeded in a 12-well plate with 50 ng/mL of hM-CSF (Peprotech, 300-25). On day 6, tumor Ge518_PTPRZ1-KI cells were stained with the CellTrace™ Far Red Cell Proliferation Kit (Invitrogen, C34564) and 25’000 cells were added to the wells, followed 2h later by the addition of anti-PTPRZ1 RNA 471_28z CAR-T cells or Mock EP control cells, at a 3:1 E:T ratio. After 72h, cells were collected, and tumor and macrophage death was evaluated by flow cytometry using a LIVE/DEAD™ Fixable Violet Dead Cell Stain Kit (Invitrogen, L34964). An anti-CD14^PE^ antibody (BioLegend, 367104) was used to detect macrophages.

### CAR expression over time and tonic signaling

Jurkat triple reporter cells were electroporated with anti-PTPRZ1 CAR mRNA, following a previously described protocol. After ON incubation, Jurkat cells were cultured for 24h alone (to evaluate tonic signaling) or co-cultured at a 1:1 E:T ratio with Ge518_PTPRZ1-KI cells (to evaluate specific signaling) or antigen-negative Ge1302 cells (to evaluate non-specific signaling). Jurkat activation was evaluated by measuring the expression of NF-κB (CFP), NFAT (eGFP), and AP-1 (mCherry) by flow cytometry (BD FACSymphony™, BD Biosciences). CAR expression was measured at 4h, 96h and 168h after electroporation using biotinylated goat anti-human IgG F(ab’)₂ (Jackson Immunoresearch, 109-066-097) followed by streptavidin conjugated to APC (BD Biosciences, 554067).

### Phenotyping of CAR-T cells by qPCR

Total cell mRNA was extracted using the RNeasy Plus Mini Kit (Qiagen, 74136) and then converted to cDNA using PrimeScript™ RT Master Mix (Perfect Real Time, Takara, RR036A). Quantitative real-time reverse transcription-polymerase chain reaction (qPCR) was performed using the primer pairs listed in Table S1. qPCR was performed using PowerUp SYBR Green Master Mix (Applied Biosystems, A25742), and the reactions were measured using a QuantStudio 6 Flex Real-Time PCR System (Applied Biosystems, Thermo Fisher Scientific). The relative fold change in gene expression compared to the 471_IgG1H_28z CAR-T cells is represented in the graph.

### Animal experiments

Experiments with animals were performed following the rules established by the Swiss Federal Animal Protection Act and complementary laws, and under the regulations of the Institutional Animal Care and Use Committee of the cantonal veterinary office (license no. VD3717_2021). NSG (NOD scid gamma, RRID:IMSR_NM-NSG-001) male mice were purchased from the Animalerie du site d’Epalinges (University of Lausanne) and used at 8–10 weeks of age. To generate an orthotopic GBM tumor model, 25’000 Ge518_PTPRZ1-KI_Luc^+^ cells were stereotactically infused into the left striatum, 2.6 mm to the bregma on day 0. Successful tumor implantation was confirmed on day 5 by luminescence detection using an IVIS® Lumina S5 Imaging System (PerkinElmer). Animals were then randomized and received an intracranial intratumoral administration of 1×10^6^ anti-PTPRZ1 RNA CAR-T cells or Mock EP T cells in 2 μL of HBSS on day 7. Tumor growth was monitored twice a week by imaging using the IVIS system. Mice were sacrificed when they showed neurological symptoms, weight loss ≥ 15% over one week, or a poor body condition score.

### Immunofluorescence

To analyze PTPRZ1 expression in the tumor after *in vivo* experiments, several cryosections were taken from tumor-bearing mice. Immediately after death, mice were perfused with PBS1X (Gibco, 10010) to eliminate intravascular red blood cells. Mouse brains were dissected, and a slice of the brain containing the tumor was included in OCT (FSC22 blue frozen section compound, Leica Biosystems, 3801480). Sectioning was performed using the Agora histology platform and the sections were stored at −80°C. For staining, the sections were warmed to room temperature (RT), fixed with 4% PFA (Thermo Fisher Scientific, 047377.9L) and blocked with 5% FBS (Sigma, F7524-500mL), 0.5% BSA (Gibco, 15260), 0.3% Triton X-100 (Sigma, T8787-250ML), and 0.1% Sodium Azide (Sigma, 71289) in PBS1X containing MOM reagent (ReadyProbes™ Mouse-on-Mouse IgG Blocking Solution, Invitrogen, R37621) at RT. To determine antigen expression, samples were stained at RT with anti-PTPRZ1 scFv RB471 conjugated to rabbit Fc and mouse anti-human HLA-ABC (eBioscience, 16-9983-85) antibodies diluted in the blocking solution. The slices were washed with PBS1X 0.3% Triton X-100, and secondary goat anti-rabbit IgG^AF555^ (Invitrogen, A21428) and goat anti-mouse IgG^AF488^ (Invitrogen, A11001) antibodies were added at RT in the dark. To evaluate human T-cell infiltration in tumors, samples were stained with anti-human HLA-ABC and anti-human CD3^PE^ (BioLegend, 317308) at RT in the dark. The slides were subsequently washed thrice with PBS1X 0.3% Triton X-100 and once with PBS1X. Slides were then mounted with Fluoromount-G mounting medium containing DAPI (Invitrogen, 00-4959-52), dried ON at 4°C and read on a Zeiss Slide Scanner (Zeiss). Images were analyzed using QuPath v0.4.3 software (RRID:SCR_018257).

### Statistical analysis

GraphPad Prism 10 software was used for all the statistical analyses (RRID:SCR_002798). All experiments were performed at least twice unless indicated otherwise. Data are shown as the mean ± standard deviation. Two-tailed unpaired Student’s t-tests or one-way ANOVA tests were used to compare the different experimental groups. For the post-tests, we used Tukey’s and Dunnett’s multiple comparison tests. In the case of the *in vivo* experiments, comparison of the differences in overall survival was made with Kaplan-Meier survival analysis, and p-values were determined using the log-rank test. In all experiments, the significance level was set at (*) p<0.05, (**) p<0.01, (***) p<0.001, and (****) p<0.0001.

### Data availability

All the data generated in this study are available in the article itself or in Supplementary Figures/Tables. The raw data are available upon request from the corresponding author.

## Results

### PTPRZ1 is overexpressed in GBM

We previously identified PTPRZ1 as a relevant tumor-associated cell surface protein in GBM through peptidome screening of GBM samples from 32 patients, and subsequent protein expression validation in 250 primary or recurrent GBM samples (12). Analysis of bulk mRNA expression in healthy tissues showed that PTPRZ1 expression was mainly restricted to the brain/spinal cord and, to a minor degree, to the skin (Figure 1A). However, PTPRZ1 was significantly more expressed in GBM vs. non-tumor samples in bulk RNA sequencing data (TCGA and Rembrandt datasets, Figure 1B). In addition, single-cell RNA sequencing data showed that PTPRZ1 was specifically expressed by neoplastic cells and not by the tumor microenvironment (Figure 1C). Interestingly, PTPRZ1 was highly expressed in cells with high oligodendrocyte precursor cell (OPC)-like scores (Figures 1D and 1E), suggesting that progenitor-like tumor cells express high levels of PTPRZ1. PTPRZ1 expression level was not associated with survival in GBM patients (Supplementary Figure 1A), but interestingly, showed high expression in glioma patients independent of different clinicopathologic features such as grade, 1p19q codeletion, IDH mutation, MGMT methylation status, and GBM subtypes (Supplementary Figure 1B-F).

**Figure 1:**
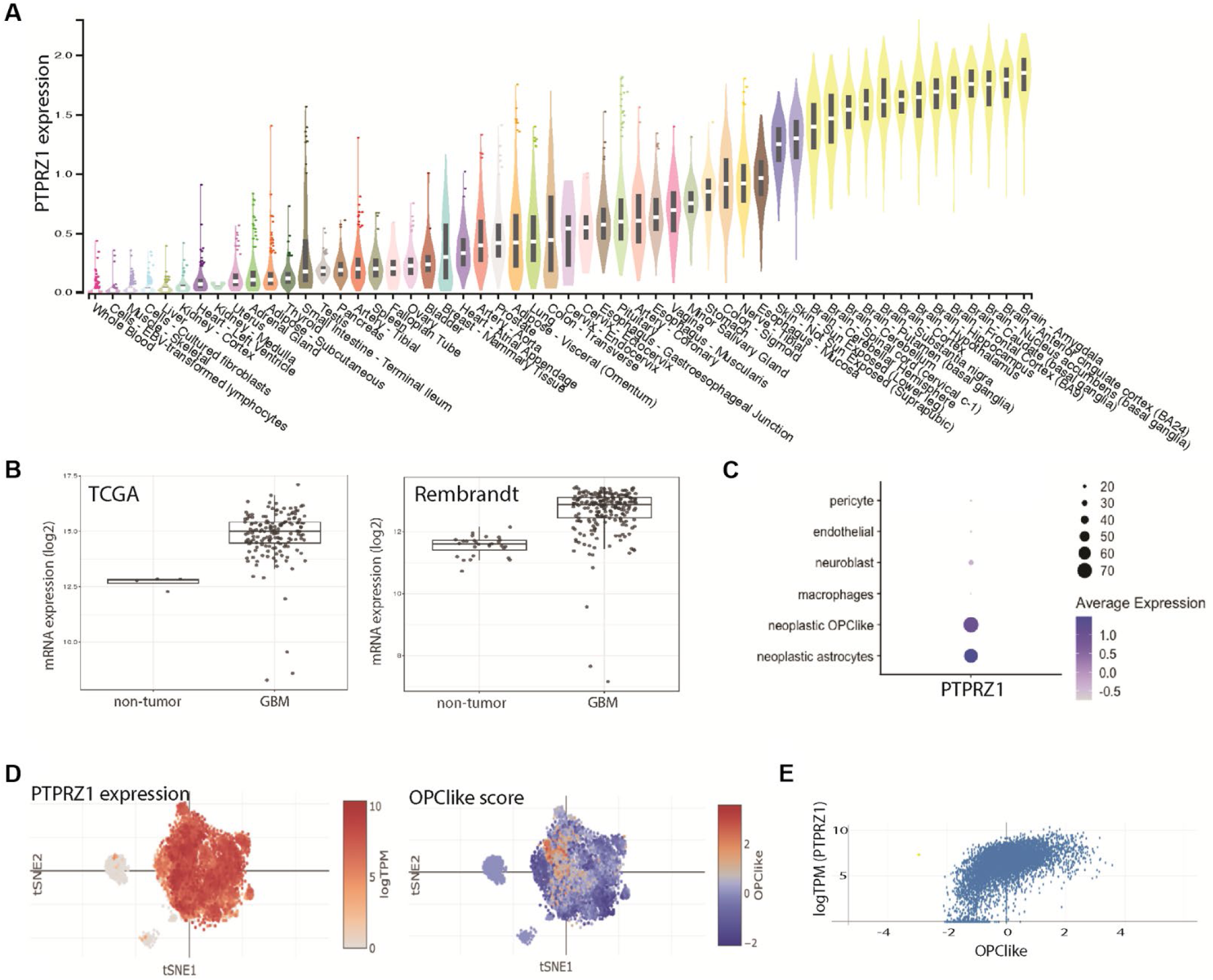
Expression of PTPRZ1 in healthy and tumor tissues at the mRNA level. **(A)** Bulk mRNA expression of PTPRZ1 in normal tissues according to GTEx RNASeq data (GTEx Analysis Release V8, dbGaP Accession phs000424.v8.p2). Data processing and normalization: expression values are shown in log10 (TPM+1); TPM= transcripts per million. Violin plots and boxplots show values of median, 25^th^ and 75^th^ percentiles; points are outliers above or below 1.5 times the interquartile range. **(B)** Boxplots showing log2 expression (median, quartiles and outliers) of PTPRZ1 in GBM and non-tumor samples in TCGA RNAseq (left) and Rembrandt microarray (right) datasets. Both comparisons between GBM and non-tumor are significant with p<0.001. **(C)** scRNAseq analysis from Yuan et al., 2018 (25). Dotplot showing average expression (color intensity) of PTPRZ1 in different cell type clusters (pericyte, endothelial, macrophages and neoplastic astrocyte-like, neuroblast-like and OPC-like). The size of the dot indicates the percentage of antigen-expressing cells in each cluster. **(D)** tSNE plot of scRNAseq from Neftel et al., 2019 (24). All malignant cells (larger cluster in the center of the projection) express PTPRZ1 (left panel). Cells with high OPC scores are colored orange/red (right panel). **(E)** Scatter plot showing the correlation between PTPRZ1 expression and OPC score.

### Generation of anti-PTPRZ1 RNA CAR-T cells with *in vitro* killing capacities

Using a human phage display library, we obtained six scFvs (RB469, RB470, RB471, RB473, RB474, and RB476) capable of binding to the α-carbonic anhydrase or fibronectin type III extracellular domains of PTPRZ1. Five of the scFvs specifically recognized the α-carbonic anhydrase domain, and one, RB470, recognized the fibronectin type III domain (Figure 2A). The affinity of scFv for the α-carbonic anhydrase PTPRZ1 domain was measured by SPR (Figure 2B). Whereas the protein did not bind to RB470 given its specificity for the Fibronectin type III PTPRZ1 domain, the KD of the other scFv ranged from 57.4 nM for RRB473 to 106.6 nM for RB476. The affinity of RB469 scFv could not be assessed because of its production at a too-diluted concentration (<10 µg/mL). Irrelevant carbonic anhydrase II protein was used as a negative control and did not bind to any scFv (not shown). We used these scFvs to generate 2^nd^ generation CAR constructs containing human CD3ζ combined with human CD28 or 4-1BB intracellular domains (Figure 2C). All CAR constructs were produced as mRNA and used to produce RNA CAR-T cells. We first generated the 4-1BB/CD3ζ (BBz) version of CAR-T cells, which showed high CAR molecule expression, except CAR 474_BBz (Figure 2D). To test the efficacy of the anti-PTPRZ1 CAR, we overexpressed PTPRZ1 in a Ge518 patient-derived GBM cell line. The six anti-PTPRZ1 scFv genes were used to assess PTPRZ1 expression in Ge518_PTPRZ1 KI cells. We observed high binding of scFvs RB469, 471, 473 and 476 and lower binding of scFvs RB470 and 474 (Figure 2E). Anti-PTPRZ1 RNA CAR-T cells were then used in an *in vitro* killing assay of the Ge518_PTPRZ1-KI cell line, and CAR-T cells 470_BBz, 471_BBz, and 476_BBz showed a high percentage of specific lysis (Figure 2F). Similar results were observed when IFN-γ was measured in the supernatant of the killing assay. Only CAR-T cells with a high percentage of tumor lysis also showed significant IFN-γ production (Figure 2G). RB470, 471 and 476 scFvs were selected for further experiments.

**Figure 2:**
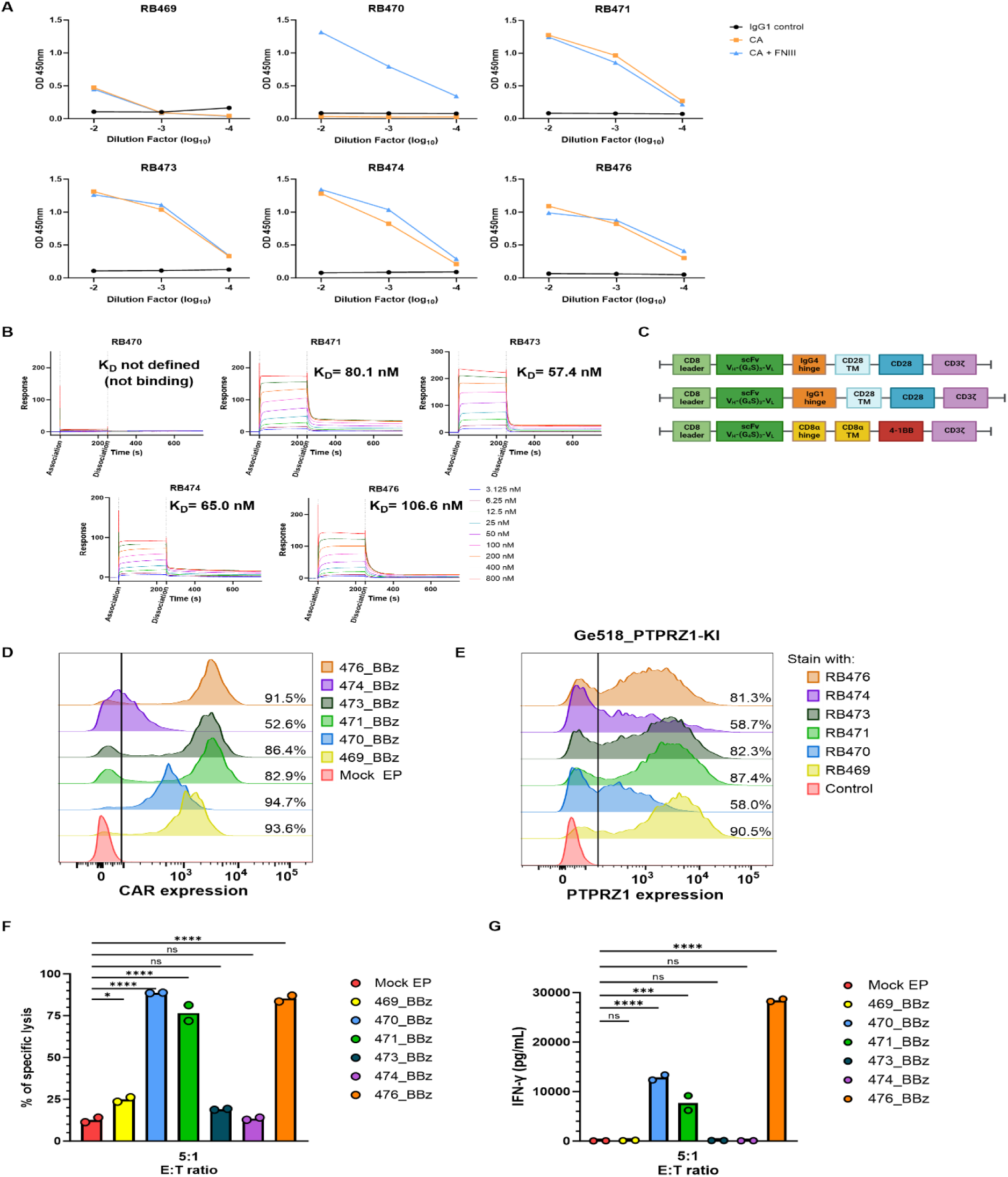
Isolation of PTPRZ1-specific scFv and generation of anti-PTPRZ1 RNA CAR-T cells. **(A)** Recognition of α-carbonic anhydrase (CA) or Fibronectin type III (FNIII) extracellular domains of PTPRZ1 by different anti-PTPRZ1 scFvs was assessed by ELISA **(B)** Affinity of the six scFv was measured by SPR: the scFv were bound to the chip and the CA protein domain was run at different concentrations. Dissociation constant (K_D_) values are shown. **(C)** Diagrams of the CAR constructs. Created with Biorender.com (RRID:SCR_018361). **(D)** CAR expression on RNA CAR-T cells expressing the different anti-PTPRZ1 scFv bearing the 4-1BB costimulatory domain (BBz). **(E)** Recognition of Ge518_PTPRZ1-KI by anti-PTPRZ1 scFv (each scFv at 2.5 µg/mL). Control is the anti-rabbit IgG^AF488^ secondary antibody alone. **(F)** *In vitro* killing of Ge518_PTPRZ1-KI by the six different anti-PTPRZ1 CAR-T cells as measured by flow cytometry. E:T ratio: 5:1. **(G)** IFN-γ secretion by the six different anti-PTPRZ1 CAR-T cells after 72h incubation with Ge518_PTPRZ1-KI tumor cells. E:T ratio: 5:1.

### 471_28z CAR-T cells show best tumor-killing capacity

Given the relatively short lifespan of RNA CAR expression, we reasoned that the CD28 co-stimulatory domain would be superior to 4-1BB because CD28 generates a faster response, with an increased proliferation rate and higher levels of Th_1_ cytokine secretion (32,33). Therefore, we generated 470_28z, 471_28z, and 476_28z CAR-T cells and compared their killing capacity (Figure 3A, left). The 471_28z CAR-T cells displayed the highest killing capacity at all ratios tested and, importantly, were still efficient at an E:T ratio of 0.5:1. This result was consistent with IFN-γ secretion by CAR-T cells (Figure 3A, right). A larger profile of cytokine secretion was measured using the 13-plex human CD8/NK panel (Figure 3B). 471_28z CAR-T cells showed a 100-times increase in TNF-α and IL-2 secretion over Mock EP T cells and more than 10 times increase in IFN-γ, IL-17A, and IL-6, whereas secretion of Granzyme B, soluble FasL, Granzyme A was in–5-10-fold range (Figure 3B). Cytokine secretion by 476_28z or 470_28z CAR-T cells compared to Mock EP cells was minimal. We then used an alternative measure of CAR-T cell cytotoxic activity through an imaging-based system (Incucyte) that allowed for the evaluation of tumor cell growth inhibition. We confirmed the better anti-tumor activity of 471_28z CAR, which was able to control tumor growth even at the lowest E:T ratio tested, whereas 470_28z CAR-T cells only controlled tumor growth temporarily at a 3:1 ratio, and 476_28z CAR-T cells were unable to control tumor growth at any E:T ratio (Figure 3C). To directly compare the CD28 and 4-1BB costimulatory domains, we tested 471_28z and 471_BBz CAR-T cells against the Ge518_PTPRZ1-KI cells. The 471_28z variant displayed slightly increased cytotoxic activity and a tendency toward higher IFN-γ secretion than the 471_BBz variant (Figure 3D). However, 471_28z CAR-T cells secreted significantly higher levels of IL-2 and TNF-α (Figure 3E). 471_28z CAR-T cells also showed similar levels of inhibition of Ge518_PTPRZ1-KI cell growth compared to 471_BBz when measured using the Incucyte assay, with the 471_28z variant being more efficient at lower E:T ratios (Figure 3F). We repeated these experiments in a second GBM cell line with forced PTPRZ1 expression (Ge1302_PTPRZ1-KI). Again, 471_28z CAR-T cells showed the highest level of tumor killing and IFN-γ production compared with 470_28z and 476_28z CAR-T cells (Supplementary Figure 2A). Moreover, 471_28z CAR-T cells showed high cytotoxic activity and a tendency toward higher IFN-γ secretion than 471_BBz CAR-T cells (Supplementary Figure 2B). These results prompted us to select 471_28z CAR for further characterization.

**Figure 3:**
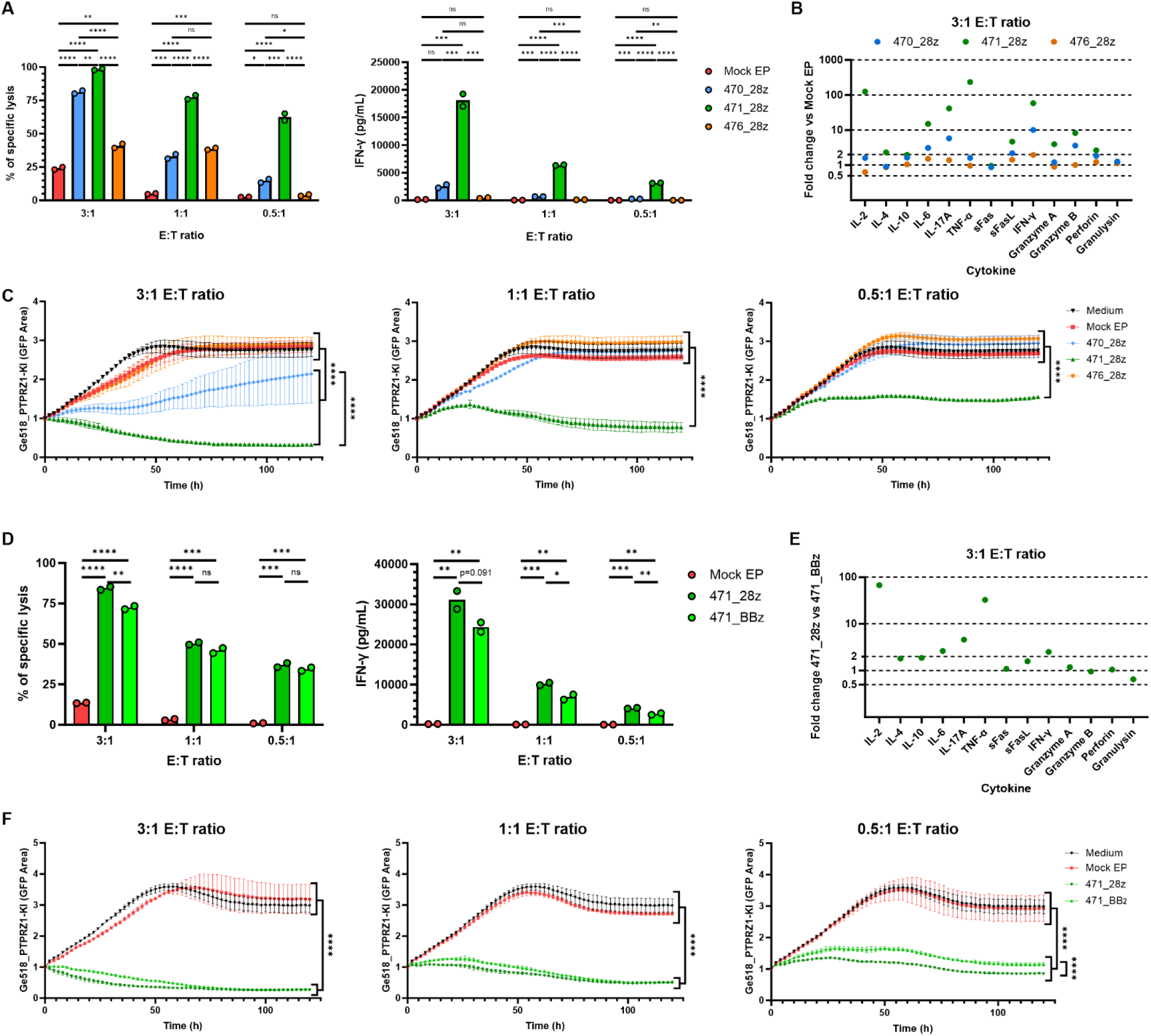
Characterization of anti-PTPRZ1 CAR-T cells in the 28z format. **(A)** *In vitro* flow cytometry killing of Ge518_PTPRZ1-KI target cells by anti-PTPRZ1 CAR-T cells bearing the 470_28z, 471_28z or 476_28z CARs at different E:T ratios (left). IFN-γ secretion by the anti-PTPRZ1 CAR-T cells from the killing assay at 72h (right). **(B)** Soluble factors secreted by the 470_28z, 471_28z or 476_28z CAR-T cells after 72h incubation with Ge518_PTPRZ1 cells at an E:T ratio of 3:1. Data are shown as fold change over levels obtained with Mock EP T cells. **(C)** *In vitro* tumor growth inhibition measure by Incucyte of Ge518_PTPRZ1-KI incubated with the 470_28z, 471_28z or 476_28z CAR-T cells or Mock EP T cells at different E:T ratios. **(D)** *In vitro* killing of Ge518_PTPRZ1-KI after incubation with BBz and 28z variants of 471 CAR-T cells at different E:T ratios (left). IFN-γ secretion by the anti-PTPRZ1 CAR-T cells from the killing assay at 72h (right). **(E)** Soluble factors secreted by BBz and 28z variants of 471 CAR-T cells after 72h incubation with Ge518_PTPRZ1-KI cells at an E:T ratio of 3:1. **(F)** *In vitro* tumor growth inhibition measured by Incucyte of Ge518_PTPRZ1-KI incubated with BBz and 28z variants of 471 CAR-T cells at different E:T ratios. Mock EP T cells were used as control throughout.

### Hinge size does not affect 471_28z CAR-T cell anti-tumor activity

Because hinge size can affect CAR-T cell activity (32–34), we evaluated whether the exchange of the 471_28z short human IgG4 hinge (12 aa) for a longer human IgG1 hinge (236 aa) influenced CAR-T cell killing capacity. Co-incubation of Ge518_PTPRZ1-KI cells with 471_IgG4H_28z or 471_IgG1H_28z CAR-T cells showed similar killing activity (Figure 4A, left panel), whereas IFN-γ secretion slightly increased in 471_IgG4H_28z CAR-T cells (Figure 4A, right panel). Additionally, qPCR analysis of genes involved in T cell activation or suppression, as well as memory and effector phenotypes, did not show major differences between CAR-T cells using the short or long hinge after incubation with Ge518_PTPRZ1-KI cells. Most of the measured genes showed a fold change between the short and long hinges in the range of 0.5 to 2 times, with *TCF7* and *TNF* showing greater variation (Figure 4B). When comparing the secretion profile of cytokines and other effector molecules by 471_IgG4H_28z and 471_IgG1H_28z CAR-T cells after incubation with Ge518_PTPRZ1-KI cells, we observed that, except for IL-2, which was secreted at higher levels by 471_IgGH4_28z CAR-T cells, the other molecules showed very comparable values (Figure 4C). When evaluating the ability of both CAR-T cells to inhibit tumor growth using the Incucyte assay, we observed a better anti-tumor activity of 471_IgG4H_28z CAR-T cells at reduced E:T ratios (Figure 4D). Finally, we observed higher cytotoxic activity and a tendency for higher IFN-γ secretion by IgG4H CAR-T cells when comparing short- and long-hinge variants against Ge1302_PTPRZ1-KI cells (Supplementary Figure 2C). Since the use of the longer IgG1 hinge shows similar or inferior results compared to the use of the smaller IgG4 hinge, and considering that a small gene load usually correlates with better transfection efficacy, we decided to use CAR-T cells bearing the short hinge for further characterization.

**Figure 4:**
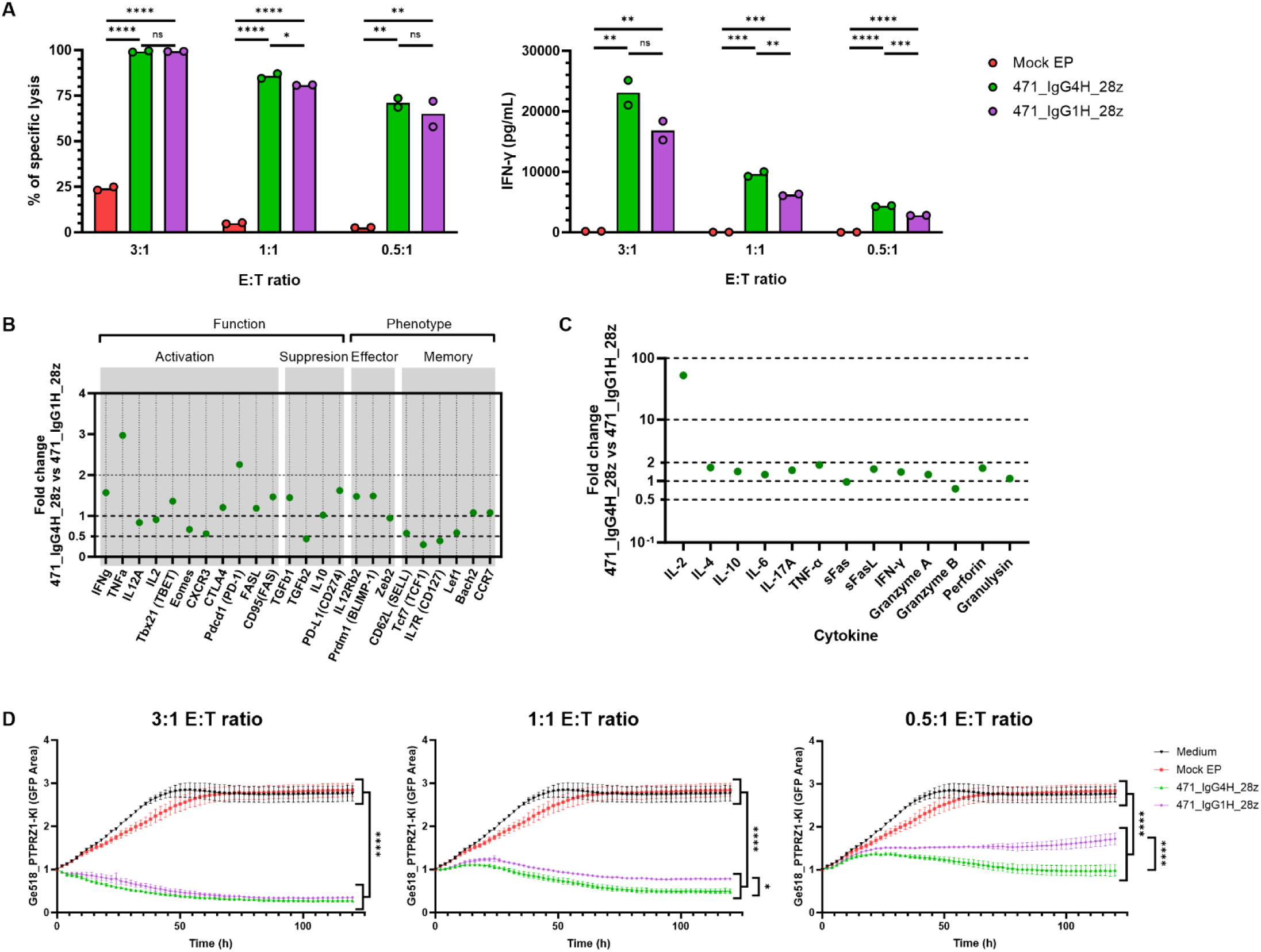
Comparison of 471_28z CAR-T cells bearing a short or long hinge. **(A)** *In vitro* killing of Ge518_PTPRZ1-KI with 471_28z CAR-T cells bearing a short (IgG4H) or a long (IgG1H) hinge at different E:T ratios (left). IFN-γ secretion by CAR-T cells from the killing assay at 72h (right). **(B)** qPCR analysis of genes involved in T cell function (activation and suppression) and phenotype (effector and memory) in 471_28z CAR-T cells bearing a short (IgG4H) or long (IgG1H) hinge after 72h incubatio n with Ge518_PTPRZ1-KI cells at an E:T ratio of 2:1. **(C)** Soluble factors secreted by 471_28z CAR-T cells bearing a short (IgG4H) or long (IgG1H) hinge after 72h incubation with Ge518_PTPRZ1-KI cells at an E:T ratio of 3:1. **(D)** *In vitro* tumor growth inhibition measured by Incucyte of Ge518_PTPRZ1-KI incubated with 471_28z CAR-T cells bearing a short (IgG4H) or long (IgG1H) hinge at various E:T ratios. Mock EP T cells are used as control throughout.

### 471_28z CAR-T cells elicit a bystander effect on antigen-negative tumor cells

To assess CAR-T cell specificity, we generated an antigen-negative control cell line by knocking out PTPRZ1 from the Ge518 cell line (Ge518_PTPRZ1-KO). 471_28z CAR-T cells specifically killed antigen-positive (Ge518_PTPRZ1-KI) cells while sparing antigen-negative (Ge518_PTPRZ1-KO) cells (Figure 5A, left), with corresponding IFN-γ secretion profiles (Figure 5A, right). These results were confirmed by a tumor cell growth inhibition assay (Figure 5B). Then, to test for potential bystander cytotoxicity of anti-PTPRZ1 CAR-T cells, antigen-positive (Ge518_PTPRZ1-KI) or antigen-negative (Ge518_PTPRZ1-KO) cells transduced with different reporter proteins were mixed in a 1:1 ratio and incubated with CAR-T or Mock EP T cells. Under these conditions, 471_28z CAR-T cells were able to kill Ge518_PTPRZ1-KI cells as well as Ge518_PTPRZ1-KO cells (Figure 5C, left) with significant IFN-γ secretion (Figure 5C, right). Similarly, 471_28z CAR-T cells could control the growth of antigen-positive and antigen-negative cells (Figure 5D). We then analyzed the secretion profile of cytokines and effector molecules by 471_28z CAR-T cells when incubated with Ge518_PTPRZ1-KI, Ge518_PTPRZ1-KO, or a combination of both cell lines. We observed that, compared to Mock EP T cells, 471_28z CAR-T cells did not show a significant increase in cytokine secretion after incubation with Ge518_PTPRZ1-KO cells, except for a 2 to 5 times increase in IFN-γ, Granzyme B, IL-6 and IL-17A secretion, which might reflect tonic signaling (Figure 5E). However, incubation with Ge518_PTPRZ1-KI or the Ge518_PTPRZ1-KI/Ge518_PTPRZ1-KO mix induced a significant increase in several cytokines, with >1000 fold increase as compared to Mock EP T cells seen for IL-2 and TNF-α, >100 fold increase for Granzyme B and IFN-γ and > 10 fold increase for IL-17A, IL-6, and soluble FasL (Figure 5E, left: relative expression, right: absolute measure of IL-2, TNF-α, IFN-γ and Granzyme B). The amount of cytokines secreted by 471_28z CAR-T cells after incubation with Ge518_PTPRZ1-KI or the combination of KI and KO cells was similar. We further showed in a transwell experiment that bystander killing was at least partly mediated by the secretion of soluble molecules, potentially cytokines (Supplementary Figure 3A-C). Importantly, bystander cytotoxic activity of 471_28z CAR-T cells was not observed when macrophages were used as target cells (Supplementary Figure 3D-G).

**Figure 5:**
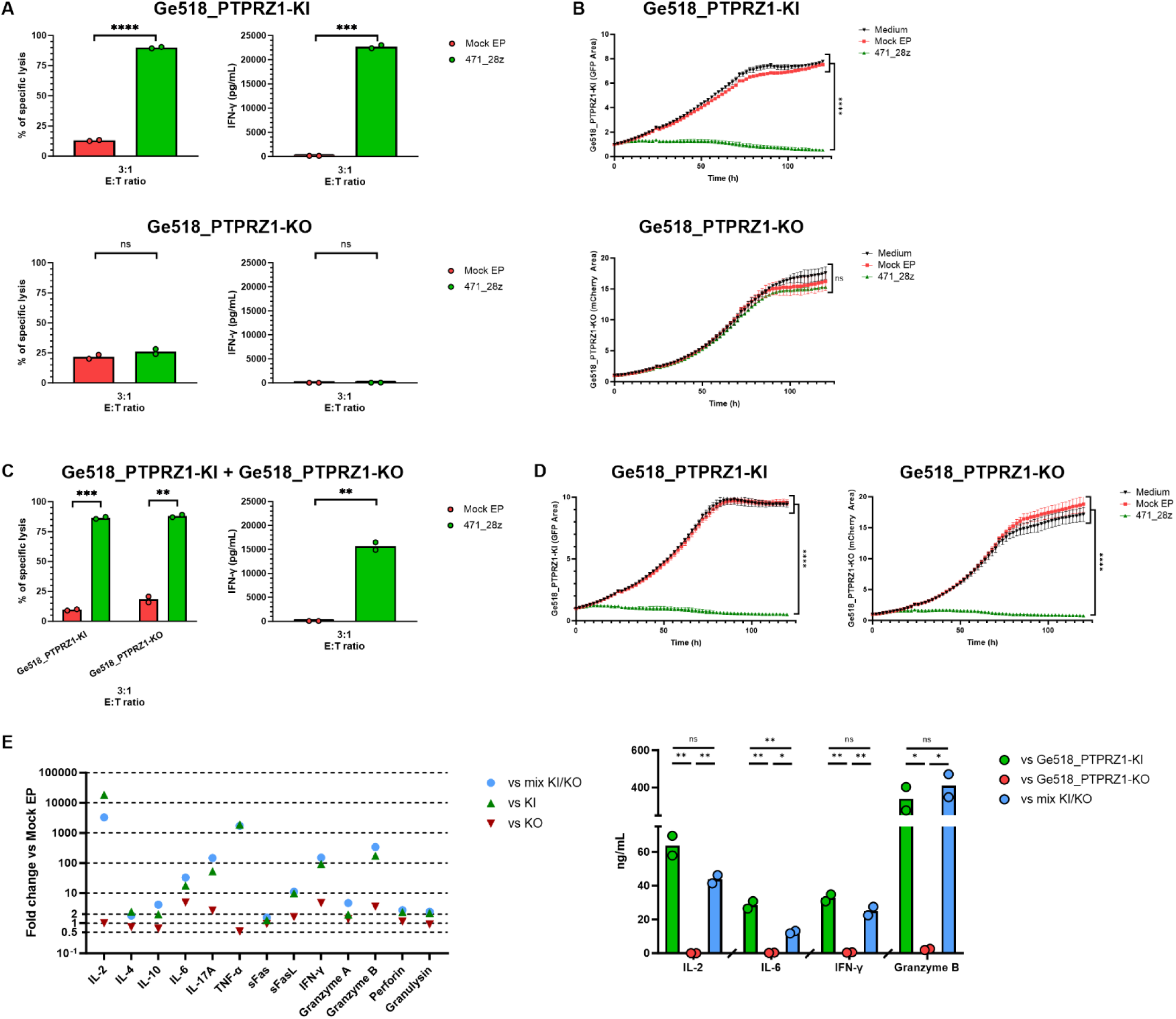
471_28z CAR-T cells show specific as well as bystander killing capacity. **(A)** Tumor cell killing, IFN-γ secretion and **(B)** tumor growth inhibition after coincubation of 471_28z CAR-T cells at 3:1 E:T ratio with Ge518_PTPRZ1-KI or Ge518_PTPRZ1-KO cells. **(C)** Tumor cell killing, IFN-γ secretion and **(D)** tumor growth inhibition after coincubation of 471_28z CAR-T cells at 3:1 E:T ratio with a 1:1 mix of Ge518_PTPRZ1-KI (GFP) and Ge518_PTPRZ1-KO (mCherry) cells. Each tumor cell population was identified by the expression of a different reported gene. **(E)** Soluble factors secreted by 471_28z CAR-T cells after co-incubation with Ge518_PTPRZ1-KI, Ge518_PTPRZ1-KO or a 1:1 mix of both cells at 72h. Left: levels are expressed as fold change vs. secretion by Mock EP T cells; right: concentration of selected soluble factors secreted by 471_28z CAR-T cells (IL-2, IL-6, IFN-γ and Granzyme B).

### 471_28z CAR-T cells display low tonic signaling

To assess the spontaneous activation of CAR-T cells, we used Jurkat cells bearing reporters for NF-κB (CFP), NFAT (eGFP), and AP-1 (mCherry), which are transcription factors downstream of T-cell activation (28). Jurkat triple reporter cells were electroporated with 470_28z and 471_28z CARs (Supplementary Figure 4A), and the expression of NF-κB, NFAT, and AP-1 was measured in the absence of coincubation with tumor cells. Compared to Mock EP T cells, a low increase in the expression of NFAT and AP-1 was observed in both CAR-expressing Jurkat cell lines (Supplementary Figure 4B). The higher values observed for 471_28z CAR were probably due to the higher expression of CAR (Supplementary Figure 4A). An increase in the transcription factor reporter expression, especially NFAT and AP-1, was observed when CAR-expressing Jurkat cells were incubated for 24h with antigen-positive Ge518_PTPRZ1-KI cells, but not with antigen-negative Ge1302 cells (Supplementary Figure 4B), supporting the specificity of both CAR molecules.

### 471_28z CAR-T cells mantain an effector memory phenotype after tumor killing

To characterize the phenotype of 471_28z CAR-T cells, we evaluated memory cell populations using CD45RO/CD62L markers and activation/exhaustion using TIM3 and PD-1 markers in both CD8 (CD3^+^CD8^+^) and CD4 (CD3^+^CD8^-^) T cell subpopulations (Supplementary Figure 5A). We compared the cell phenotype before freezing (pre-cryo), ∼16h after thawing (post-thawing), and after a 72h-killing assay against Ge518_PTPRZ1-KI tumor cells (post-killing). We did not observe significant differences between the CAR-T cell phenotypes before freezing and after thawing (Supplementary Figure 5B-E). However, we detected a relative decrease in the CD8/CD4 ratio in CAR-T cells compared to Mock EP T cells after the 72h killing assay as compared to post-thawing (Figure 6A). In addition, whereas CD4 T cells from both 471_28z and Mock EP T cells showed a high proportion of T_CM_ memory cells after thawing, a significant decrease in T_CM_ cells and a concomitant increase in the T_EM_ population were observed in CD4 CAR-T cells after killing, which was not observed in Mock EP T cells (Figure 6B, upper panels). We observed a similar decrease in T_CM_ CD8 + T cells in CAR-T cells compared to Mock EP cells, with a concomitant increase in T_EFF_ cells and a tendency to increase T_EM_ cells (Figure 6B, lower panels). The change from a central memory to a more effector phenotype was also evidenced by a decrease in CD127^+^ cells in both T-cell subpopulations (Figure 6C). Regarding exhaustion markers, whereas 471_28z CAR-T cells mostly expressed PD-1 after thawing, the proportion of PD-1^+^ CAR-T cells decreased after the killing assay compared to Mock EP T cells. There was a concomitant but insignificant increase in TIM3^+^ and PD-1^+^TIM3^+^ in CAR-T cells compared to Mock EP cells, which was probably due to the heterogeneity in the percentage of expressing cells among the three different experiments (Figure 6D, upper panels for CD4 and lower panels for CD8 T cells).

**Figure 6:**
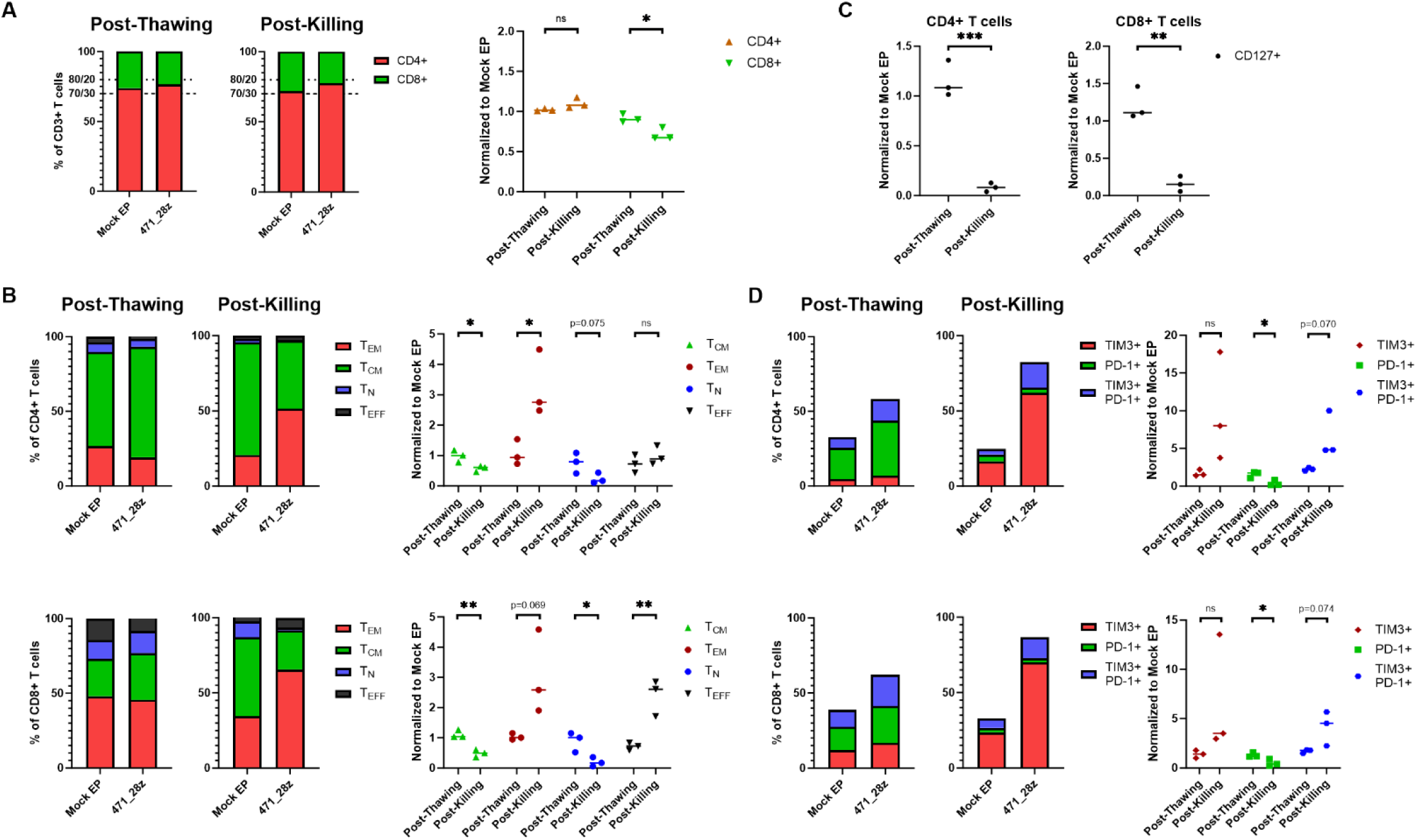
Expression of effector/memory and activation/exhaustion markers by CAR-T cells in post-thawing vs. post-killing samples. **(A)** Proportion of CD4^+^ and CD8^+^ T cells in the CD3^+^ cell population in Mock EP and 471_28z CAR-T cells. Right panel: expression in 471_28z CAR-T relative to that in Mock EP cells **(B)** Proportion of central memory (T_CM_), effector memory (T_EM_), effector (T_EFF_) and naïve (T_N_) populations in CD4^+^ (top row) and CD8^+^ (bottom row) T cells in 471_28z CAR-T and Mock EP T cells. Right panel: expression in 471_28z CAR-T relative to that in Mock EP T cells. **(C)** Comparison of CD127 expression in 471_28z CAR-T relative to that in Mock EP T cells in CD4^+^ (left) and CD8^+^ (right) T cells. **(D)** Percentage of PD-1^+^, PD-1^+^/TIM3^+^ and TIM3^+^ CD4^+^ (top row) and CD8^+^ (bottom row) T cells in 471_28z CAR-T cells and Mock EP T cells. Right panels: expression in 471_28z CAR-T relative to that in Mock EP T cells. Individual dots represent different donors tested in different experiments.

### RNA 471_28z CAR-T cells are able to delay tumor growth *in vivo*

Finally, we evaluated the anti-tumor capacity of 471_28z CAR-T cells *in vivo* using an orthotopic GBM model. 25’000 Ge518_PTPRZ1-KI_Luc^+^ cells were orthotopically injected into the left striatum of NSG mice, and the mice were treated one week later with 1×10^6^ anti-PTPRZ1 471_28z CAR-T cells, Mock EP T cells, or HBSS controls (Figure 7A). Tumor growth was followed by the luminescence of Luc^+^ cells. In contrast to control mice in which tumors grew continuously, tumor growth was arrested for approximately seven days in mice treated with 471_28z CAR-T cells, leading to a delay in tumor progression (Figure 7B and C). As a result, 471_28z CAR-T cell-treated mice showed significantly prolonged overall survival, which increased by ∼45% compared to the control groups (Figure 7D). The 471_28z CAR-T cells were well tolerated, as we did not observe any changes in animal weight after injection (Figure 7E). A second *in vivo* experiment showed a similar seven-day delay in tumor progression (Supplementary Figure 6A) and a significant increase in overall survival (Supplementary Figure 6B). Mouse brain sections showed that PTPRZ1 expression did not decrease in animals treated with 471_28z CAR-T cells, excluding antigen loss as the cause of tumor progression (Supplementary Figure 6C). Additionally, tumors of mice treated with CAR-T cells did not show a significant increase in human T-cell infiltration, probably due to transient CAR expression (Supplementary Figure 6D).

**Figure 7:**
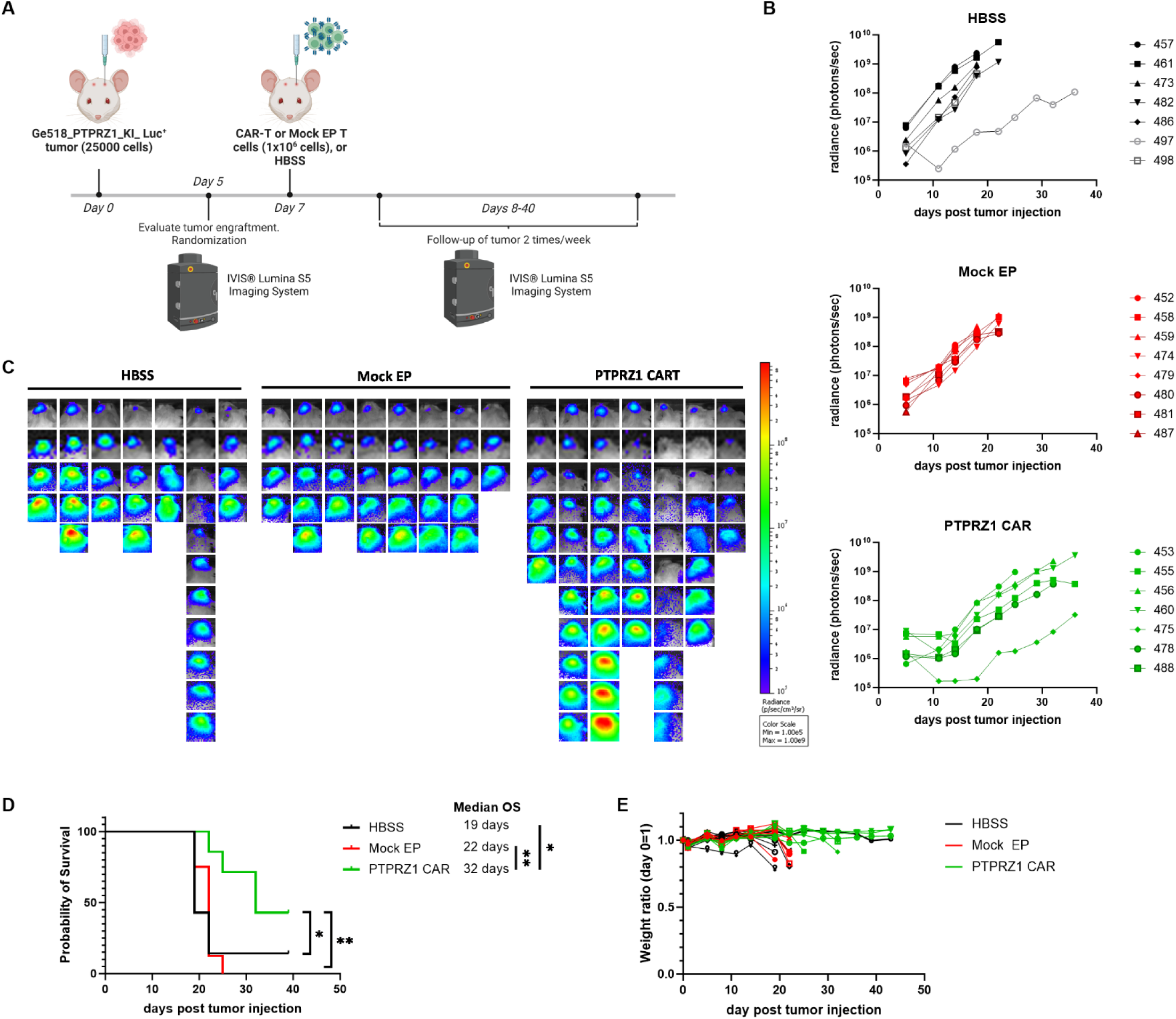
*In vivo* anti-tumor activity of anti-PTPRZ1 CAR-T cells. **(A)** Scheme showing *in vivo* experiments testing the activity of anti-PTPRZ1 CAR-T cells. Created with Biorender.com. **(B)** Tumor volume evolution was measured by luminescence in the HBSS, Mock EP and anti-PTPRZ1 CAR-T cells groups (n=6-7 mice per group). **(C)** Imaging of tumors by luminescence in all mice included in the experiment. **(D)** Overall survival in mice treated with Mock EP cells, anti-PTPRZ1 CAR-T cells or HBSS (*p=0.0383 CAR-T vs HBSS, **p=0.0018 CAR-T vs Mock EP). **(E)** Mouse weight variation during the *in vivo* experiment. Weight is expressed as a normalized ratio to each animal’s weight at day 0.

## Discussion

Immunotherapy with CAR-T cells has shown great success in hematopoietic malignancies but has not yet been successfully extended to solid tumors, including GBM. Here, we identified PTPRZ1 as an attractive target that can be considered as a tumor-associated antigen in GBM, with low expression in healthy tissues of the brain and peripheral organs. We confirmed the association of PTPRZ1 expression with glioma stem cell properties, which is consistent with other reports (16,18,35). In addition, PTPRZ1 is becoming an emerging target in GBM given its detection as a relevant tumor-associated molecule in immunoproteomics (12,36), single-cell transcriptomics (37), and tumor biology studies (16,17). Here, we report for the first time the generation of anti-PTPRZ1 CAR-T cells, which show specific anti-tumor activity against GBM cell lines both *in vitro* and *in vivo*.

In this study, we used RNA-based CAR-T cells, as opposed to conventional lentivirus-based technology. The use of *in vitro*-transcribed mRNA confers certain advantages to CAR-T cell immunotherapy (21,22). Among them is the reduction in manufacturing time and use of potentially lower doses due to high efficiency in CAR expression, which, combined with transient CAR expression, results in greater safety by avoiding excessive proliferation and accumulation of cytokines that typically lead to cytokine-release syndrome (CRS) and immune effector cell–associated neurotoxicity syndrome, the main side effects of lentivirus-based CAR-T cells (22,38,39). RNA CAR-T cells have been previously generated against other brain tumor antigens, such as GD2 or GPC2, showing high levels of cytotoxicity, increased survival in mouse tumor models, and no evidence of toxicity (40,41). Several clinical trials using RNA CAR-T cells against solid tumors or autoimmune diseases have also demonstrated the safety of this approach (42,43). The major limitation of RNA CAR-T cells is related to transient CAR expression, which prevents long-term control and might require injection of multiple doses, which might be challenging for patients with limited availability of T cells (21). Our study in mice demonstrated that a single dose of anti-PTPRZ1 CAR-T cells could temporarily arrest tumor growth and improve animal survival without showing any evidence of toxicity. Administration of several CAR-T cell doses might lead to a full antitumor effect (44) but might be limited in mice due to the side effects of multiple surgical procedures. However, in the clinical setting, recent reports have shown that multiple doses of CAR-T cells can be administered to patients with brain tumors without increasing the side effects (45).

The antitumor capacity of CAR-T cells can be optimized in different ways (33). In our case, we selected scFv with the best antitumor activity from a panel of six and evaluated the impact of hinge length and the co-stimulatory domain. Hinge length influences the establishment of an optimal immunological synapse distance and, thus, the proper functioning of CAR-T cells (33,34). In the case of the anti-PTPRZ1 CAR, we observed a superior cytotoxic activity of a short (IgG4) hinge over a long (IgG1) hinge, especially when IFN-y secretion and tumor growth inhibition measured by cell-based imaging were considered. This, combined with the advantage of a smaller gene size for CAR expression, led us to select a short hinge for the generation of anti-PTPRZ1 CAR-T cells. On the other hand, the use of the CD28 co-stimulatory domain in the anti-PTPRZ1 CAR construct showed a tendency for higher levels of cytotoxicity and cytokine secretion compared to 4-1BB, in line with the faster activation kinetics and superior Th1 cytokine profile associated with CD28 (32,33). These characteristics are best suited for RNA CAR-T cells owing to transient CAR expression, indicating that the CD28 costimulatory domain is the best option. Indeed, we verified that anti-PTPRZ1_28z CAR-T cells secreted the main pro-inflammatory cytokines/molecules associated with anti-tumor activity, such as IFN-γ, TNF-α and Granzymes. However, the fact that we also detected the secretion of pro-inflammatory cytokines involved in CRS, especially IL-6, must be considered. However, CRS is mostly observed in patients treated with lentivirus-based CAR-T cells compared to those treated with RNA CAR-T cells (42,43). In addition, it is important to note that macrophages, not CAR-T cells, are the main source of IL-6 during CRS (46). Finally, although human IL-6 was active in mice, no toxicity was detected in *in vivo* experiments.

Selection of the route of administration is another key element in maximizing the effectiveness of CAR-T cells in the case of brain tumors. Here, we opted for intracranial administration at the tumor site, which circumvents the problem of T cell migration to the brain and maximizes the use of RNA CAR-T cells for antitumor activity. Results from clinical trials support the idea that intracentral nervous system administration constitutes a safe and efficient route of delivery for the treatment of GBM with CAR-T cells (47,48).

Finally, the main characteristic of GBM is its high tumor cell heterogeneity, which is one of the major obstacles to the development of successful immunotherapies (11). Hence, the desired quality of any CAR-T cell product is its ability to indirectly kill local antigen-negative tumor cells. We were able to demonstrate this bystander effect using anti-PTPRZ1 CAR-T cells. This may be mediated by death receptor ligands, such as Fas, either as a membrane or soluble form, or by inflammatory cytokines such as IFN-γ (49), the latter constituting the main cytotoxicity mechanism associated with CD4^+^ CAR-T cells (50). Thus, it might be the mode of action of the bystander effect in our experiments, as the anti-PTPRZ1 CAR-T cell batches used in our study typically have a high proportion (70-80%) of CD4^+^ CAR-T cells. The bystander killing of antigen-negative tumor cells by anti-PTPRZ1 CAR-T cells through the secretion of inflammatory cytokines, especially IFN-γ and TNF-α, could be increased *in vivo* where cytokines can also target the local tumor stroma, increasing the anti-tumor activity of CAR-T cells (51,52). Importantly, we showed that this bystander effect might preferentially affect tumor cells, as human macrophages were not killed, suggesting the absence of direct toxicity in healthy cells.

In conclusion, we generated novel RNA CAR-T cells that target PTPRZ1, a tumor-associated antigen commonly overexpressed in GBM and considered a marker of glioma stem cells. We demonstrated that anti-PTPRZ1 CAR-T cells exerted a potent *in vitro* cytotoxic effect against GBM tumor lines, including the secretion of high levels of pro-inflammatory cytokines. Anti-PTPRZ1 CAR-T cells also exert antitumor activity *in vivo* by temporarily arresting tumor growth and increasing mouse survival. However, RNA-based CAR expression is time-limited, posing a significant challenge for its use in solid tumors. To overcome this limitation, we need to improve the therapeutic scheme by developing a multi-dose model or by combining anti-PTPRZ1 RNA CAR-T cells with other therapeutic modalities. In this regard, using anti-PTPRZ1 RNA CAR-T cells after the surgical removal of GBM might be a promising option for evaluation. The results reported here are the basis for the initiation of a clinical trial of anti-PTPRZ1 CAR-T cells, which we plan to open in the near future.

## Acknowledgments

We thank Isabelle Grandjean for managing mouse colonies in Agora (Lausanne), as well as the animal facility, in vivo imaging facility, the flow cytometry core, and the histology facility in Agora, especially to Véronique Noguet Brechbuehl for cutting the mouse brain slides. Some diagrams were created using Biorender.

## Author contributions

DMB, VD, and DM designed the experiments, collected and analyzed the data, interpreted the results, and drafted the paper. DMB, SD, SE, and LCC performed *in vitro* experiments. EM analyzed the sequencing data. MD, SB, and NW performed SPR experiments. KS, PB, and SM collected human GBM cells that were used to generate the primary GBM cell lines. DMB and SD performed *in vivo* GBM and IF studies. PH and PC performed the phage display experiments. DM designed the study, interpreted the results, and supervised and coordinated the study. DMB, PRW, VD, and DM wrote the paper with input from all authors.

