## Supplemental Data for "PTPRZ1-targeting RNA CAR-T cells exert antigen-specific and bystander antitumor activity in glioblastoma"

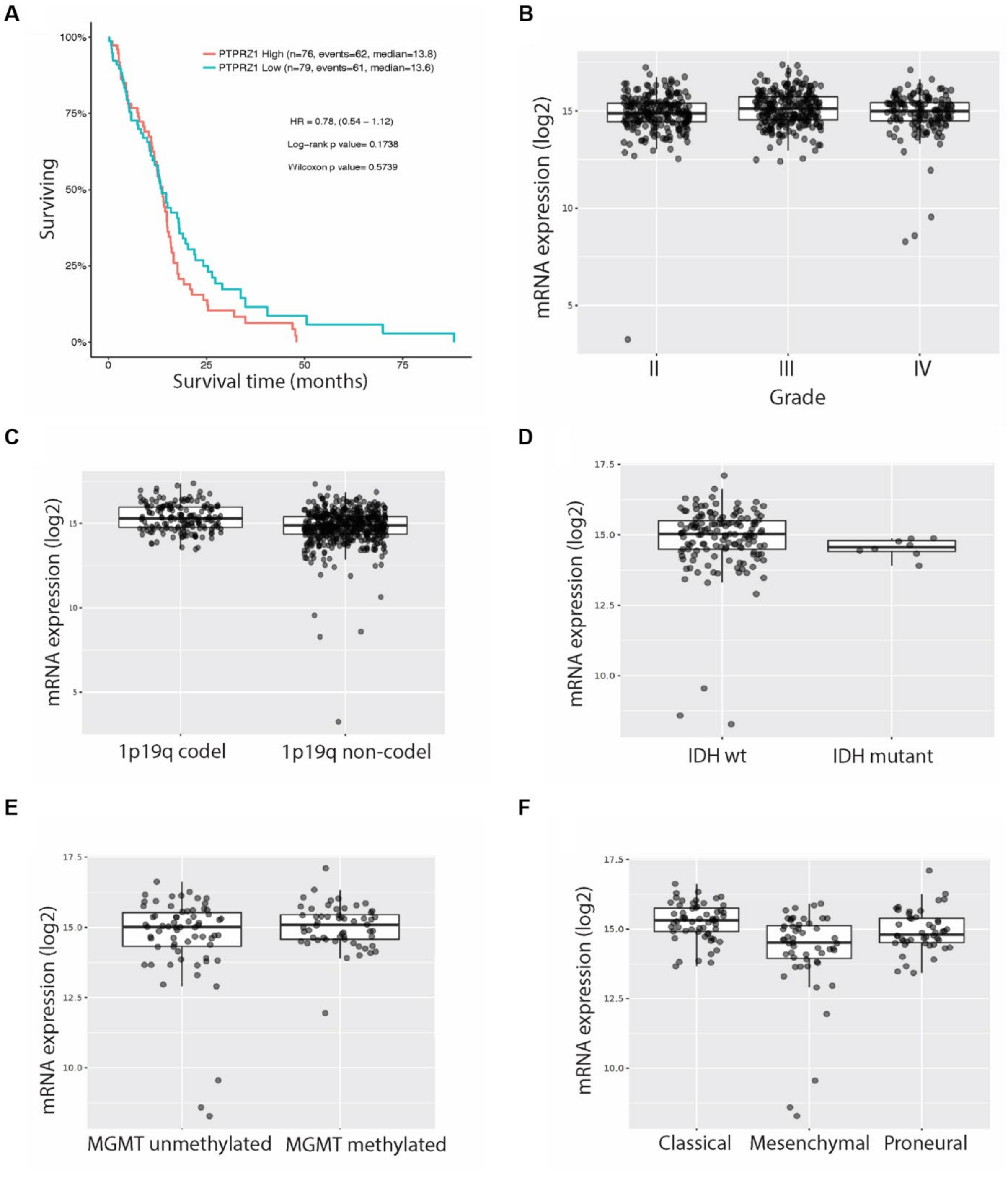


**Figure S1: mRNA level expression of PTPRZ1 according to different clinicopathologic features of GBM (A)** Kaplan–Meier survival analysis of GBM patients from TCGA RNAseq dataset was performed using the ‘survival’ package in R via Gliovis (http://gliovis.bioinfo.cnio.es). The median of PTPRZ1 expression was chosen as cut-off, log-rank p-value >0.05. Boxplots show log2 expression (median, quartiles and outliers) of PTPRZ1 in the TCGA glioma RNAseq dataset according to grade **(B)**, 1p19q codeletion **(C)**, IDH mutational status **(D)**, MGMT methylation status **(E)**, and GBM subtypes **(F)**.

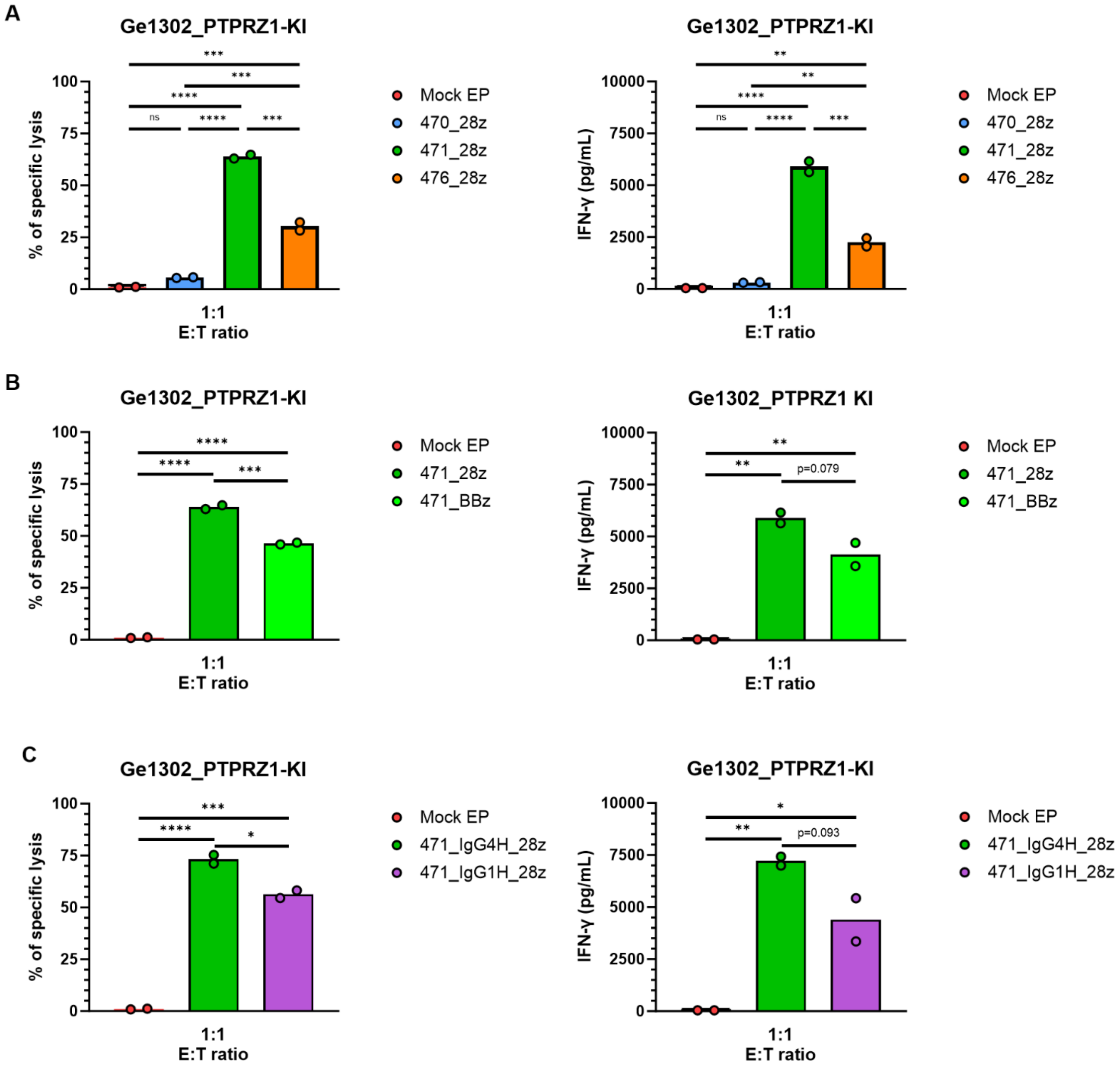


**Figure S2: Effector function of anti-PTPRZ1 CAR-T cells after incubation with the Ge1302_PTPRZ1-KI cells (A)** *In vitro* flow cytometry killing of Ge1302_PTPRZ1-KI target cells by anti-PTPRZ1 CAR-T cells bearing the 470_28z, 471_28z or 476_28z CARs at a 1:1 E:T ratio (left). IFN-γ secretion by the anti-PTPRZ1 CAR-T cells from the killing assay at 72h (right). **(B)** *In vitro* killing of Ge1302_PTPRZ1-KI incubated with BBz and 28z variants of 471 CAR-T cells at a 1:1 E:T ratio (left). IFN-γ secretion by the anti-PTPRZ1 CAR-T cells from the killing assay at 72h (right). **(C)** *In vitro* killing of Ge1302_PTPRZ1-KI with variants of 471 CAR-T cells bearing a short (IgG4H) or long (IgG1H) hinge at a 1:1 E:T ratio (left). IFN-γ secretion by CAR-T cells from the killing assay at 72h (right). Mock EP T cells were used as control throughout.

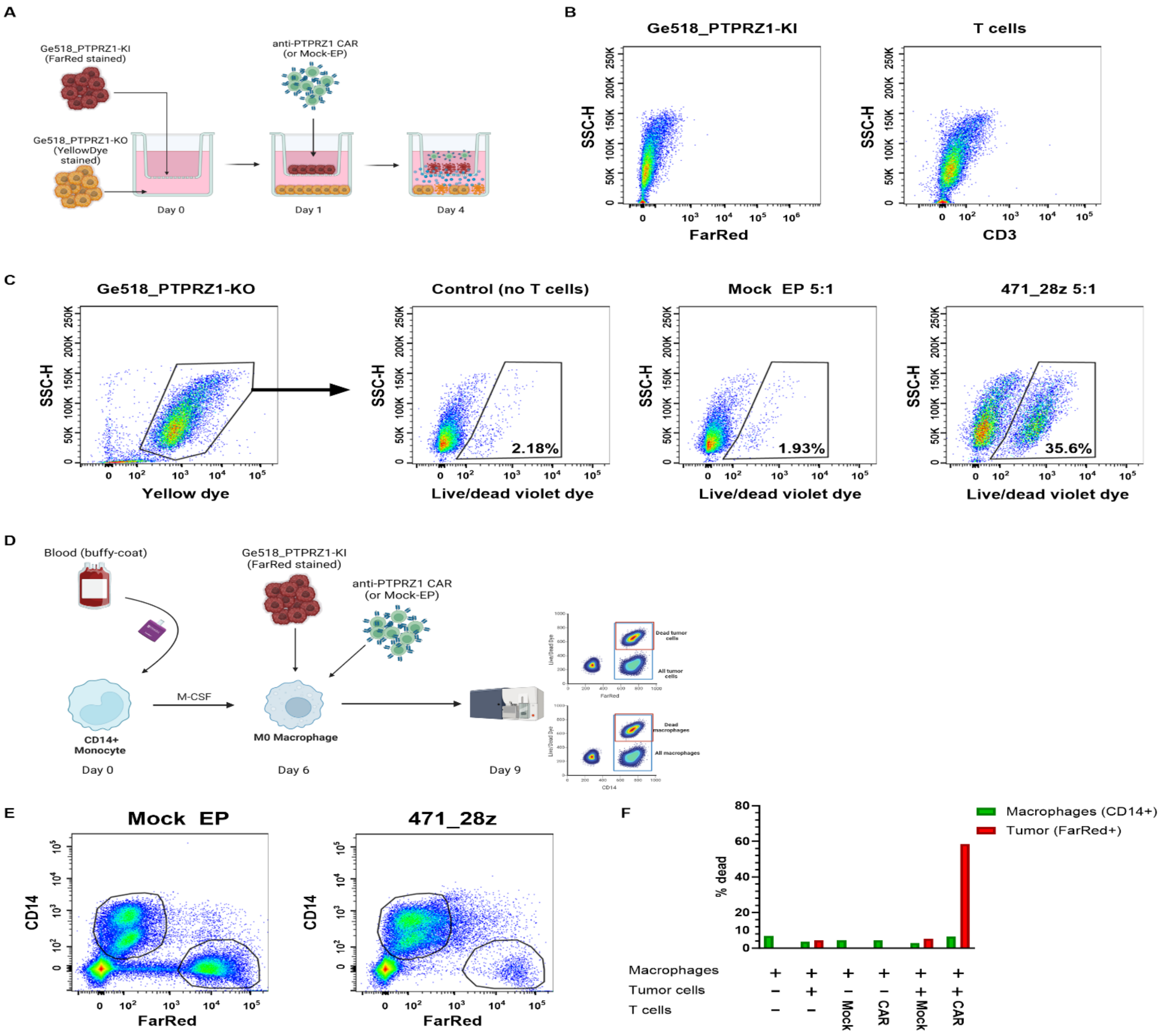


**Figure S3: Bystander killing by 471_28z CAR-T cells depends on soluble mediators and does not affect macrophages. (A)** At day 0, CellTrace Yellow labeled Ge518_PTPRZ1-KO cells were seeded at the bottom of a transwell plate and CellTrace FarRed labeled Ge518_PTPRZ1-KI cells were seeded on the top of the transwell. On Day 1, 471_28z CAR-T cells or Mock EP control cells were added on the top of the transwell at a 5:1 E:T ratio. After 72h, cells on the bottom of the well were collected and cell death was evaluated by flow cytometry. Created with Biorender.com. **(B)** Confirmation of absence of Ge518_PTPRZ1-KI cells (stained with FarRed) or T cells ($CD3^{+}$) on the bottom part of wells. **(C)** Evaluation of indirect, soluble mediator-dependent, killing of Ge518_PTPRZ1-KO cells (stained with CellTrace Yellow) by 471_28z CAR-T cells. **(D)** $CD14^{+}$ monocytes were purified from human blood at day 0 and were differentiated to macrophages by a six-day culture with M-CSF. At day 6, Ge518_PTPRZ1-KI cells (FarRed stained) were added to the macrophage culture, followed by addition of 471_28z CAR-T cells or Mock EP T cells, at a 3:1 E:T ratio. After 72h, cells were collected and death of tumor cells and macrophages was evaluated by flow cytometry. Created with Biorender.com. **(E)** Gating strategy to differentiate Ge518_PTPRZ1-KI tumor cells ($CD14^{-}$ $FarRed^{+}$) and human macrophages ($CD14^{+}$ $FarRed^{-}$) in wells containing Mock EP (left) or 471_28z CAR-T cells (right). **(F)** *In vitro* evaluation of killing of Ge518_PTPRZ1-KI or human macrophages by 471_28z CAR-T cells.

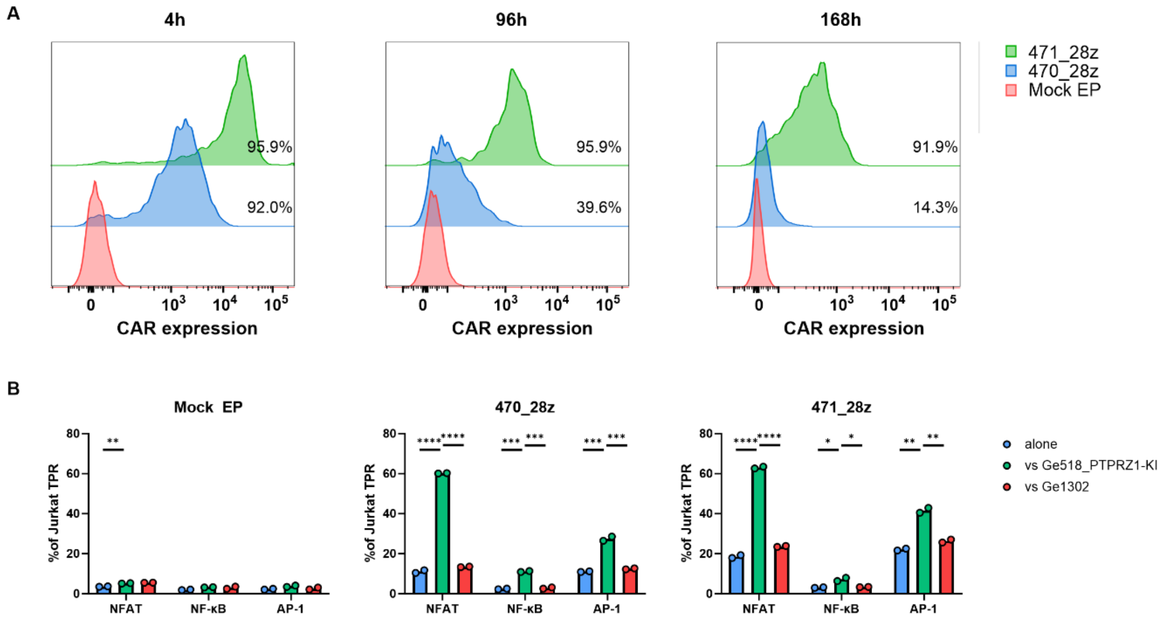


**Figure S4: Use of Jurkat triple reporter cells to assess CAR tonic signaling and specificity. (A)** Jurkat cells were electroporated with the 471_28z or 470_28z CARs and CAR expression was measured at 4h, 96h and 168h after electroporation. **(B)** Tonic signaling and specific activation of 471_28z or 470_28z CARs in Jurkat cells measured 24h after electroporation when Jurkat cells were cultured without tumor cells (alone) or incubated at a 1:1 E:T ratio with antigen-positive (Ge518_PTPRZ1-KI) or antigen-negative (Ge1302) cells.

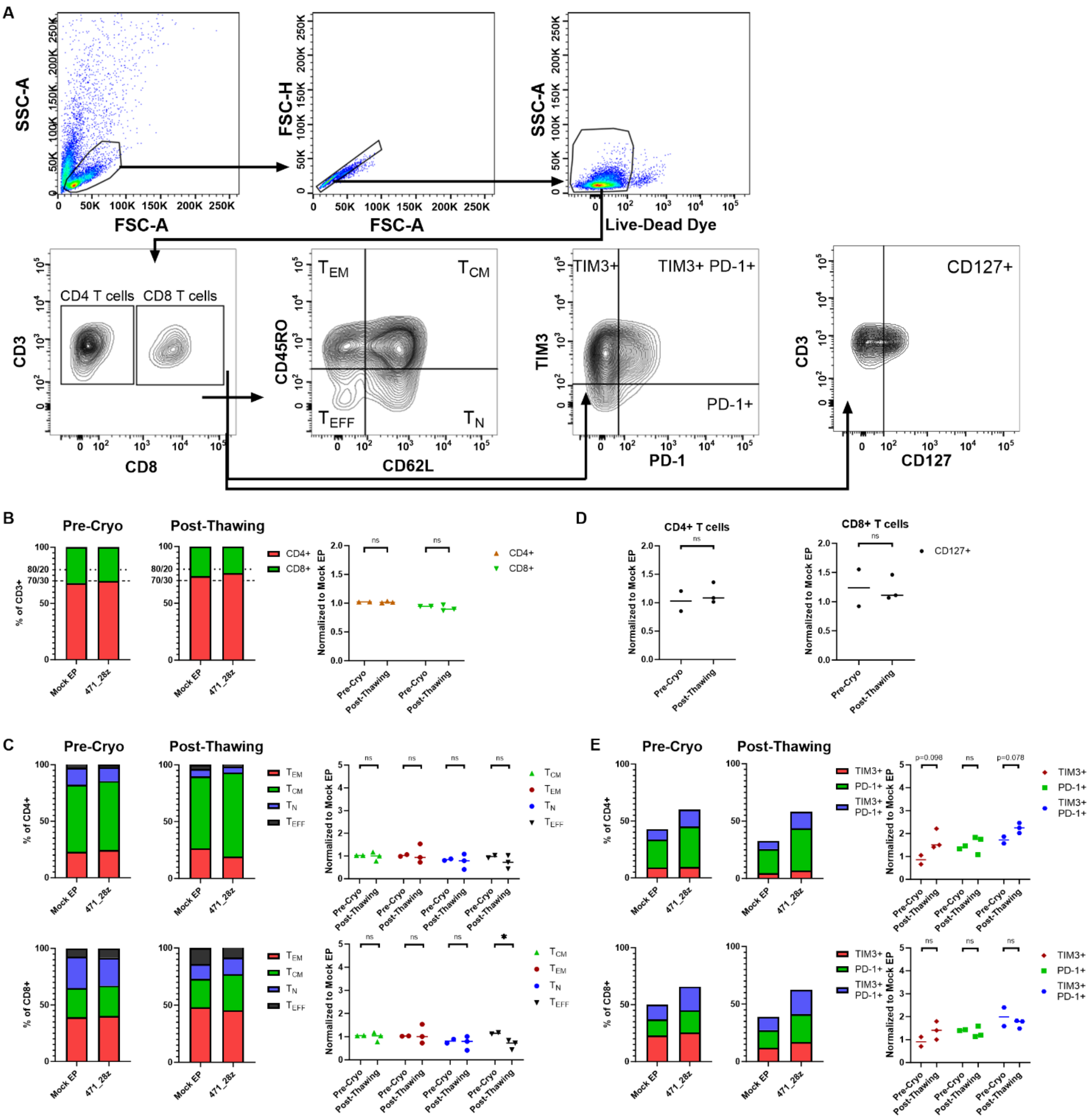


**Figure S5: Expression of effector/memory and activation/exhaustion markers by CAR-T cells in pre- vs. post-thawing samples. (A)** The gating strategy followed to determine the phenotype of CAR-T cells and Mock EP T cells is shown. **(B)** Proportion of CD4+ and CD8+ T cells in the CD3+ cell population in Mock EP and 471_28z CAR-T cells, before freezing (pre-cryo) and after thawing of the T cells. Right panel: expression in 471_28z CAR-T relative to that in Mock EP T cells. **(C)** Proportion of central memory ($T_{CM}$), effector memory ($T_{EM}$), effector ($T_{EFF}$) and naïve ($T_N$) populations in CD4+ (top row) and CD8+ (bottom row) T cells in 471_28z CAR-T cells and to Mock EP T cells, before freezing and after thawing the T cells. Right panel: expression in 471_28z CAR-T relative to that in Mock EP T cells. **(D)** Comparison of CD127 expression in 471_28z CAR-T relative to that in Mock EP T cells in CD4+ (left) and CD8+ (right) T cells, before freezing and after thawing the T cells. **(E)** Comparison of the frequency of PD-1+, PD-1+/TIM3+ and TIM3+ CD4+ (top row) and CD8+ (bottom row) T cells in 471_28z CAR-T cells and Mock EP T cells, before freezing and after thawing of the T cells. Right panel: expression in 471_28z CAR-T relative to that in Mock EP T cells. Individual dots represent different donors tested in different experiments.

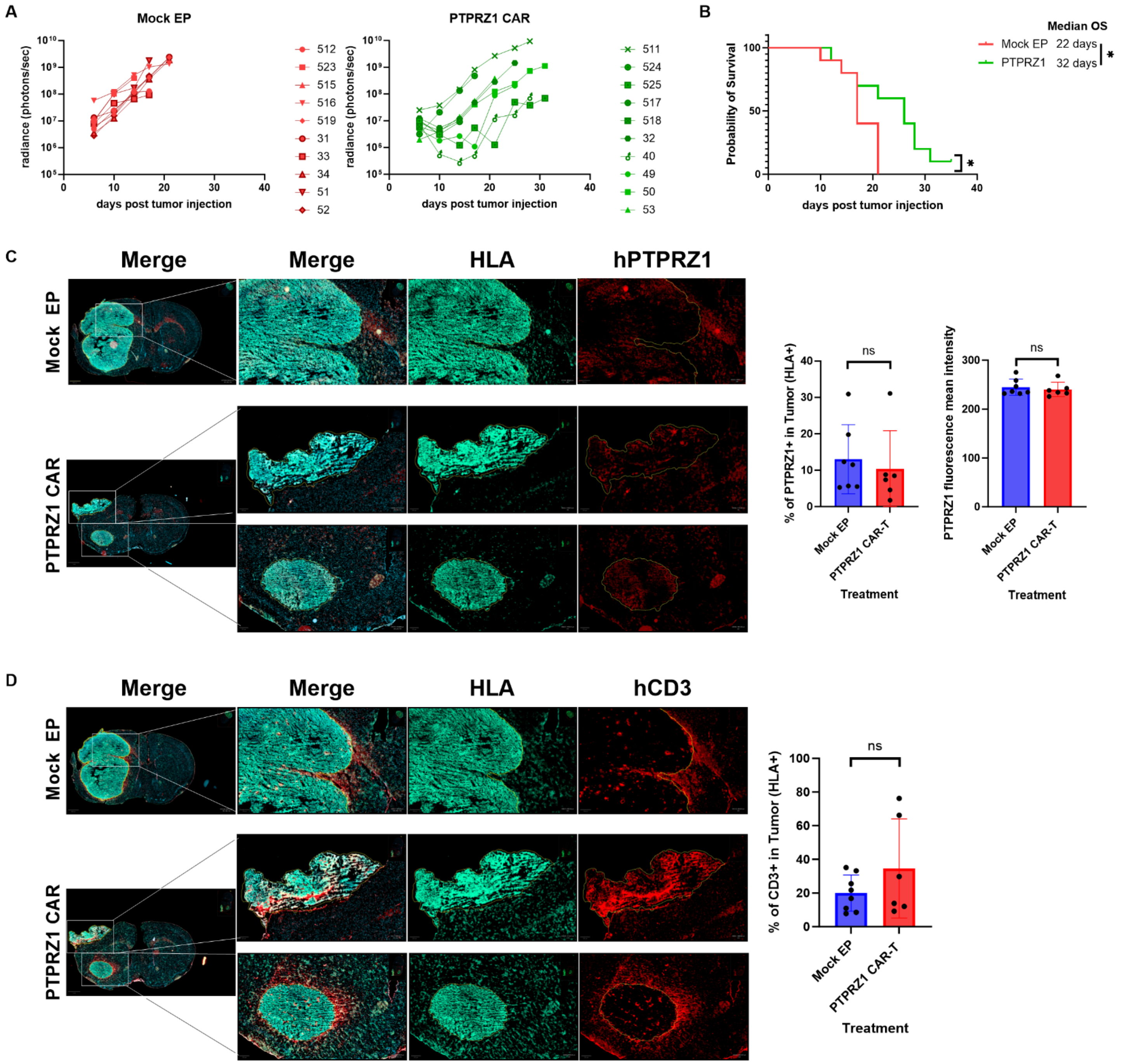


**Figure S6: *In vivo* anti-tumor activity of anti-PTPRZ1 CAR-T cells. (A)** Tumor volume evolution was measured by luminescence in the Mock EP and anti-PTPRZ1 CAR-T cells groups (n=10 mice per group). **(B)** Overall survival in mice treated with Mock EP cells or anti-PTPRZ1 CAR-T cells (*p=0.0210). **(C)** IF images of representative mouse brain samples stained with anti-human HLA-ABC (HLA) and anti-PTPRZ1 scFv RRB471 conjugated to rabbit Fc (hPTPRZ1). Panels on the right show quantification of PTPRZ1 positive area and PTPRZ1 staining fluorescence mean intensity in the tumor (defined by HLA-ABC staining). **(D)** IF images of representative mouse brain samples stained with anti-human HLA-ABC and anti-human CD3. Panel on right shows quantification of CD3 positive area in the tumor (defined by HLA-ABC staining).

**Table S1_Primer pairs for CAR-T cell phenotyping by qPCR**

| Gene name | Full name | Alternative name | Forward Primer (5'-3') | Reverse Primer (5'-3') |
|---|---|---|---|---|
| *IFNG* | interferon gamma | | TGG CTT TTC AGC TCT GCA TC | CCG CTA CAT CTG AAT GAC CTG |
| *TNF* | tumor necrosis factor | TNFA, TNFSF2, TNF-α | CCC GAG TGA CAA GCC TGT AG | TCT CAG CTC CAC GCC ATT |
| *IL12A* | interleukin 12A | | ACA GTG GAG GCC TGT TTA CCA T | GGC CAG GCA ACT CCC ATT AGT T |
| *IL12B* | interleukin 12B | | CTC TGG CAA AAC CCT GAC C | GCT TAG AAC CTC GCC TCC TT |
| *IL2* | interleukin 2 | | AGAACTCAAACCTCTGGAGGAAG | GCTGTCTCATCAGCATATTCACAC |
| *TBX21* | T-box transcription factor 21 | T-bet, TBET | ATTGCCGTGACTGCCTACCAGA | GGAATTGACAGTTGGGTCCAGG |
| *EOMES* | eomesodermin | | AAATGGGTGACCTGTGGCAAAGC | CTCCTGTCTCATCCAGTGGGAA |
| *CXCR3* | C-X-C motif chemokine receptor 3 | | ACGAGAGTGACTCGTGCTGTAC | GCAGAAAGAGGAGGCTGTAGAG |
| *CTLA4* | cytotoxic T-lymphocyte associated protein 4 | | ACGGGACTCTACATCTGCAAGG | GGAGGAAGTCAGAATCTGGGCA |
| *PDCD1* | programmed cell death 1 | CD279, PD-1, PD1 | AAGGCGCAGATCAAAGAGAGCC | CAACCACCAGGGTTTGGAACTG |
| *FASLG* | Fas ligand | CD95L, FASL, TNFSF6 | GGTTCTGGTTGCCTTGGTAGGA | CTGTGTGCATCTGGCTGGTAGA |
| *FAS* | Fas cell surface death receptor | TNFRSF6, FAS | GGACCCAGAATACCAAGTGCAG | GTTGCTGGTGAGTGTGCATTCC |
| *TGFB1* | transforming growth factor beta 1 | TGFbeta | GAG CCC TGG ACA CCA ACT AT | ATC CAC TTC CAG CCG AGG T |
| *TGFB2* | transforming growth factor beta 2 | | TTC GAT GTA ACT GAT GCT GTT C | AAT TAT TAG ATG GTA CAA AAG TGC AG |
| *IL10* | interleukin 10 | | GGT TGC CAA GCC TTG TCT GA | AGG GAG TTC ACA TGC GCC T |
| *CD274* | programmed cell death 1 ligand 1 | B7H1, PD-L1 | TGCCGACTACAAGCGAATTACTG | CTGCTTGTCCAGATGACTTCGG |
| *IL12RB2* | interleukin 12 receptor subunit beta 2 | | AGACCTCAGTGGTGTAGCAGAG | TGATGACCAGCGGTTCAGGATC |
| *PRDM1* | PR/SET domain 1 | BLIMP1 | CAGTTCCTAAGAACGCCAACAGG | GTGCTGGATTCACATAGCGCATC |
| *ZEB2* | zinc finger E-box binding homeobox 2 | | AATGCACAGAGTGTGGCAAGGC | CTGCTGATGTGCGAACTGTAGG |
| *SELL* | selectin L | CD62L, LAM1, LECAM1 | TCACAGTGTGCCTTCAGCTGCT | TCTGGTGCTGATAGAGGCTCAC |
| *TCF7* | transcription factor 7 | TCF-1 | CTGACCTCTCTGGCTTCTACTC | CAGAACCTAGCATCAAGGATGGG |
| *IL7R* | interleukin 7 receptor | CD127, IL-7R-alpha | ATCGCAGCACTCACTGACCTGT | TCAGGCACTTTACCTCCACGAG |
| *LEF1* | lymphoid enhancer binding factor 1 | TCF10 | CTACCCATCCTCACTGTCAGTC | GGATGTTCCTGTTTGACCTGAGG |

| | | | | |
|---|---|---|---|---|
| *BACH2* | BTB domain and CNC homolog 2 | | CTGCCGCAAAAGGAAACTGGAC | GGAAAGGCAGGAGAAGTTGTCC |
| *CCR7* | C-C motif chemokine receptor 7 | | CAACATCACCAGTAGCACCTGTG | TGCGGAACTTGACGCCGATGAA |
| *EEF1A1* | eukaryotic translation elongation factor 1 alpha 1 | | AGC AAA AAT GAC CCA CCA ATG | GGC CTG GAT GGT TCA GGA |
